# Quorum Sensing Regulators and Non-ribosomal Peptide Synthetases Govern Antibacterial Secretions in *Xenorhabdus szentirmaii*

**DOI:** 10.1101/2025.02.03.636359

**Authors:** Ritisha Dey, Domonique Olivia Valle, Abhijit Chakraborty, Kimberly A Mayer, Jagadeesh Uppala, Anish Chakraborty, Shama Mirza, Steven Forst, Troy Skwor, Madhusudan Dey

**Author notes:** **Corresponding author:** Madhusudan Dey (Professor,).

## Abstract

The decades-long gap in antibiotic discovery has led to a significant health crisis due to antimicrobial resistance (AMR). The bacterial genus *Xenorhabdus*, which forms symbiotic relationships with the soil nematode *Steinernema*, are known to secrete a variety of antimicrobial compounds with potential effectiveness against AMR. These antimicrobial compounds are primarily bio-synthesized by non-ribosomal peptide synthetases (NRPS) and polyketide synthetase (PKS) genes. In this study, we report that *X. szentirmaii* produces high levels of antibiotic activity during the stationary phase against diverse bacteria including known antibiotic resistant pathogens. It possesses 17 operons to encode predicted NRPS and PKS enzymes, designated as *ste1* through *ste17*. The *ste15-ste16* and *ste17* operons are predicted to produce the known antibiotics Pax peptide and Fabclavine, respectively. Additionally, the newly identified operons *ste3*, *ste4*, *ste5*, *ste8, ste9*, and *ste14* consist of single genes, each containing two or more *NRPS* genes. The *ste13* operon harbors two *NRPS* genes, while the ste7 and ste12 operons contain three *NRPS* genes each. Further, RNA-seq analysis showed that *lsrF* that encodes a quorum sensing autoinducer-2 (AI-2) aldolase was expressed at high levels during stationary phase. These findings provide evidence that *X. szentirmaii* uses quorum sensing (QS) to synchronize the expression of multiple NRPS and PKS enzymes responsible for synthesizing various antimicrobial compounds. This study underscores the potential to leverage these regulatory insights for maximizing commercial applications of novel antibiotics combating AMR, as well as broader industrial uses.

## Introduction

Antimicrobial resistance (AMR) is a major health crisis with a growing number of bacteria and fungi becoming resistant to current antibiotics [1]. Approximately 4.95 million people die each year due to AMR infections[2]. Resistance mechanisms in bacteria and fungi involve reducing drug uptake, altering the drug’s target, inactivating the drug, and actively expelling the drug[3]. This rapid rise and prevalence of AMR underscores the urgent need to discover new antibiotics from natural sources with new modes of action and new chemistries. While significant efforts have been made to identify new antibiotics from natural resources, such as the well-studied bacterial genus *Streptomyces* [4], a promising new untapped natural resource of antibiotics lies within the Gram-negative Xenorhabdus bacteria, belonging to the order of Enterobacterales [5–7].

Many of the antimicrobial compounds used in medicine today are produced by non-ribosomal peptide synthetases (NRPS) and polyketide synthetase (PKS) genes[8–12]. NRPSs also produce other secondary metabolites (SMs) such a siderophores, pigments, toxins, and immunosuppressive compounds[8, 9]. Studies reveal that NRPSs are modular enzymes of three core domains consisting of an adenylation (A) domain, a peptidyl carrier protein transfer (PCP or P) domain and a condensation (C) domain [13, 14]. The A-domain performs the selection and activation of a specific amino acid. The PCP-domain shuttles the amino acid or peptide between A and C-domains facilitated by an enzyme phosphopantetheinyl transferase encoded by a conserved gene *ngrA* [15]. Therefore, inactivation of *ngrA* abolishes the synthesis of all NRPS-derived antimicrobials [16]. The C-domain accepts the amino acid and catalyzes the formation of a peptide bond. The number of peptide bonds in a non-ribosomal peptide (NRP) depends on the number and order of the NRPS modules[17]. The modular nature of NRPSs make them flexible to produce a wide range of structurally diverse antibiotics.

Bacterial species of the genus *Xenorhabdus* form species-specific mutualistic associations by colonizing the intestines of the soil-dwelling entomopathogenic Steinernema nematodes (EPNs)[5–7]. These EPNs with the *Xenorhabdus* bacteria invade insect larvae through natural openings (e.g., mouth and anus), migrate to the insect midgut, perforate the intestinal wall and enter the hemocoel where they release their *Xenorhabdus* symbionts [8]. The released *Xenorhabdus* bacteria multiply within the insect larvae and employ a dual strategy to utilize insects as a natural resource by (I) producing diverse toxins to kill the insects and (II) secreting numerous antibiotics to suppress growth of competing microbes. Both EPNs and *Xenorhabdus* utilize the nutrient resources of the insect cadaver to support their growth and reproduction.

*X. nematophila* is the most extensively studied species of *Xenorhabdus*. It contains multiple NRPS gene clusters that encode several small peptide compounds [18, 19]. These compounds are composed of between 2-10 amino acids with a variety of modifications, including D-isomerization and cyclization[9]. One of these antibiotics, called odilorhabdin, interacts with both a specific site on the 16S rRNA of bacterial ribosome and the anticodon loop of the A-site tRNA, thus inhibiting the ribosome-mediated cellular protein synthesis[20]. Additionally, odilorhabdin has been shown to reduce bacterial infections in animal models[20]. It is noteworthy that odilorhabdin binds to a novel site on the ribosomal 16S rRNA molecule that has not been targeted by any current antibiotics, suggesting that odilorhabdin is a new type of ribosome inhibitor. While *X. nematophila* has been studied extensively, the less well studied *X. szentirmaii* was found to produce higher levels of antimicrobial activity against a variety of microbial species including animal, plant, and fungal pathogens[16]. Antibiotics of *X. szentirmaii* were also found to be active against other species of *Xenorhabdus*[16]. This makes *X. szentirmaii* an excellent source for the discovery and isolation of novel antimicrobial compounds that have potential utility in medicine and agriculture.

Tobias et al. (2017) performed a comprehensive genomic and metabolomic data of *Xenorhabdus* strains, including the draft genome sequences of *X. szentirmaii*. Their study catalogued the NRPS-containing gene clusters in *Xenorhabdus*, including five such clusters in the *X. szentirmaii* genome [21]. Additionally, they fractionated cell-free liquid cultures of *X. szentirmaii* by HPLC followed by high-resolution electrospray ionization mass spectrometry (MS). This analysis led to the identification of five compounds (i.e., Xenematide, GameX peptide, Rhabdopeptide, Xenoamicin, Szentiamide) predicted to be synthesized by NRPS clusters. It should be noted that they did not identify the compound Fabclavine, although the related NRPS gene cluster was predicted. Conversely, the compound Xenematide was identified [22], but not its corresponding gene <u>clusters. Thus,</u> the relationship between the biosynthetic gene clusters (BCGs) in the *X. szentirmaii* genome and its antibiotic production remains to be fully elucidated.

Antibiotics are a type of secondary metabolite typically produced by bacteria in planktonic culture during the stationary phase[23, 24]. These secondary metabolites confer competitive advantages by producing antimicrobial compounds (e.g., NRPs), influencing microbial survival through the synthesis of pigments and toxins (e.g., terpenes) and contributing to biofilm formation (e.g., phenazines)[25]. They also play important roles in nutrient acquisition[26] and quorum sensing[27]. Quorum sensing (QS), also known as density sensing, regulates a variety of cellular behavior, including bacterial luminescence, spore formation and biofilm formation, and antimicrobial drug resistance[28, 29]. Consequently, bacterial QS system is a promising therapeutic target for antibacterial strategies.

The QS system generally operates on three fundamental principles. First, members of the bacterial community produce signaling molecule known as autoinducers (AIs). Ais, at low cell densities, diffuse away, resulting in concentrations too low to be detected. However, at high cell densities, the cumulative production of AIs creates a localized high concentration, allowing detection and triggering a response[30]. Second, AIs are recognized by receptors located in the cytoplasm or embedded in the cell membrane. Third, detection of AIs not only activates the expression of genes required for cooperative behavior but also stimulates additional AI production[31]. This feed-forward autoinduction loop likely facilitates synchronized activity within the population and control a variety of bacterial behavior. However, the regulatory mechanism linking quorum sensing to microbial resistance and antimicrobial production is still unclear.

The antimicrobial compounds secreted by *Xenorhabdus* species have yet to achieve commercial utilization due to several technical challenges, such as insufficient knowledge of their molecular biochemistry, difficulties in large-scale purification, and the high cost [32]. To identify and characterize the antimicrobial compounds produced by *X. szentirmaii,* understanding the fundamental regulatory mechanisms governing antibiotic production by its NRPS and PKS genes is essential. In this study, we adopted a global bioinformatics approach to identify the potential biosynthetic gene clusters in the *X. szentirmaii* genome associated with antibiotic production. This involved performing NCBI BLAST searches using the PCP domain, followed by systematic analysis of the identified biosynthetic gene cluster. Through RNA-sequencing and subsequent mutational analyses, we identified novel operons with optimal expression in the stationary phase, which are associated with antimicrobial production and regulated by QS mechanisms.

## Materials and Methods

### Bacterial strains

The following strains were used in this study: *Xenorhabdus szentirmaii strain DSM16338 (nematode Steinernema Rerum), Xenorhabdus nematophila strain ATCC19061 (nematode Steinernema carpocapsae), Staphylococcus saprophyticus, Escherichia coli* DH5α, a clinical ESBL-producing uropathogenic *E. coli* MWDL6 strain[33], methicillin-resistant *Staphylococcus aureus* ATCC 43300, *Enterococcus faecium* ATCC 7171, *Staphylococcus epidermidis, Klebsiella pneumoniae* subsp. *pneumoniae* ATCC 33495, *Neisseria flava, Streptococcus pneumoniae* ATCC 6305, *Acinetobacter baumannii* ATCC 19606, and *Candida albicans* (ATCC 14053).

### Overlay assay

Five μL culture of *X. szentirmaii* (OD_600_ = 1) was spotted on an LB agar plate. The plate was placed in a 30°C incubator overnight. The cells were then killed by chloroform vapor. One ml of tester strain (OD_600_ = 1) was mixed into 5 ml of soft agar medium (LB plus 0.7% agar) and overlaid onto the LB agar plate containing the killed bacteria. The plate was placed into a 37°C incubator overnight. For overlay assay with the cell-free supernatant: 5 μL of cell-free supernatant was spotted and dried on the LB-agar plate before overlaying with the tester strain.

### Cross-streak analysis

Placed 30 µl of a diluted *X. szentirmaii* culture in the middle of a 150mm Mueller Hinton (MH) plate and incubate for 48 h at 30°C. On the back of the plates, draw radiating lines 0.5 cm from the *X. szentirmaii* colony. Overnight cultures of each test strain were diluted to a 0.5 MacFarland and then streaked alone its designated line. Growth inhibition was measured post-24 h and recorded. As a negative control to ensure neighboring bacteria did not inhibit each other, 30 µl of sterile phosphate buffered saline was added to a separate MH plate and test strains plated in the same orientation. No inhibition was evident between any of the strains (data not shown).

### Plasmid construction

Two primers (forward primer with a Kpn1 site and reverse primer with a Bamh1 site) were used to amplify the upstream ∼500 bases of our gene of interest (*ste2, ngrA* or *lsrF*) using the primers of interest (see **Table 3**). Both pKNOCK plasmid and PCR products were digested with the Kpn1-BamH1. The digested pKNOCK vector was then ligated with the PCR product to create various pKNOCK-yfg (<u>y</u>our favorite <u>g</u>ene) plasmids and transformed into *E. coli* S17 competent cells.

### Bacterial gene disruption by conjugation

The S17 cells containing the plasmid pKNOCK-yfg was mixed with wild type *X. szentirmaii* cells in a 1:1 ratio (OD_600_ = 1). The cells were plated on an LB agar plate and placed in a 30°C incubator overnight. The cells were then streaked on an LB agar plate containing ampicillin (50 μg/ml) and chloramphenicol (20μg/ml). The gene disruption was confirmed by PCR using an upstream primer (**Table 2** for primers) and a reverse primer designed from the chloramphenicol gene.

### RNA isolation and reverse-transcriptase (RT)-PCR

Four single colonies of the bacterium *X. szentirmaii* were grown individually in a liquid medium in the presence of ampicillin (50 mg/ml). Cells were harvested at the log phase (OD_600_=-0.5) and stationary phase (OD_600_=1.5). Total RNA was prepared by standard protocol using PureLink RNA isolation kit (Invitrogen, USA). Using 2 μg of total RNA, cDNA was prepared by random primers (NEB, USA, cat # S1330S) and NEB cDNA synthesis kit (NEB, USA, cat # E6300S). cDNA was then amplified by gene-specific primers (*ngrA, ste2 or LsrF*, see **Table 2**).

### Library preparation, sequencing and RNA-seq analysis

RNA quality, determined using a NanoDrop 1000 spectrophotometer (Thermo Fisher Scientific), Bioanalyzer 2100 (Agilent), and Qubit fluorometer (Thermo Fisher Scientific), were: 260/280 ratio 2.0-2.1, 260/230 ratio 2.0-2.3, and RIN>9. Sequencing libraries were prepared from 1μg of total RNA from each sample using Illumina Stranded Total RNA Prep with Ribo-Zero Plus or Ribo-Zero Plus Microbiome (M-GL-02148 v1.0) and IDT for Illumina DNA/RNA UD INdexes (Illumina, 20026121). Prepared libraries were then sequenced using an Illumina MiSeq, with paired end reads of 150 bp.

The raw sequencing data were pre-processed using the FASTP program[34] to ensure quality control, trim adapters, and prune reads. The resulting FASTQ files were aligned with the *X. szentirmaii* genome (NIBV01000001.1 and NIBV01000002.1) using the STAR (Spliced Transcripts Alignment to a Reference) aligner tool[35] with default parameters and the “alignIntronMax” parameter set to 1. Across all replicates, the uniquely mapped read fraction exceeded 70%, with absolute mapped reads ranging from 1.57 to 3.61 million. We used the gene-wise counts from the STAR to perform differential gene expression analysis via the DESeq2 program (v1.38.3)[36]. In this process, genes with a cumulative count of less than 10 across replicates were excluded. The remaining genes were analyzed using DESeq2 default parameters. A gene was deemed differentially expressed if its adjusted p-value was below 0.05 and it exhibited an absolute log2-fold change exceeding 1. To conduct principal component analysis (PCA), we utilized variance-stabilized gene expression values obtained via the “vst” function and derived the principal components using the “plotPCA” function of DESeq2 program.

### Genome analysis

The *X. szentirmaii* genome sequences (accession numbers NIBV01000001 and NIBV01000002) were retrieved from NCBI database. The NCBI BLAST tool was used to BLAST the PCP domain sequences (RDPIEIELCT TFEQILSVKR VGIHDDFFEL GGHSLLAVKL VNHLKKAFGT ELSVALLAQY STVERLGEII RENKE) against the *X. szentirmaii* genome (Max target sequences 5000, BLOSUM62, expected threshold 0.05) DNA sequence was analyzed by SnapGene software, protein sequences were analyzed and peptide sequences were predicted by using the website nrps.igs.umaryland.edu).

## Results

### 1. The X. szentirmaii genome contains 17 operons that encode NRPS module

NRPSs are modular enzymes of three core domains consisting of an adenylation (A) domain (∼500 amino acids), a peptidyl carrier protein transfer (PCP or T) domain (∼70-90 amino acids) and a condensation (C) domain (∼350 amino acids) (**Fig 1A**) [13, 14]. To become functional, the PCP-domain of NRPS and PKS transitions from their inactive apo-forms to the functional halo-forms through the covalent attachment of the 4′-phosphopantetheine (P-pant) to a conserved serine residue. This reaction is catalyzed by the phosphopantetheinyl transferase (PPTase). In *Xenorhabdus spp*, the PPTase is encoded by the gene *ngrA* [15]. The *ngrA*-PPTase is a conserved protein among *Xenorhabdus* species and is predicted to adopt a pseudo-2-fold architecture [37] (**Fig 1B**) like the *Bacillus subtilis* Sfp-type PPTase [38]. Disruption of the *ngrA* gene in *Xenorhabdus* spp. resulted in loss of antimicrobial secretion [8–11], suggesting that NRPS and PKS synthesized antimicrobials in *Xenorhabdus*. To further understand the role of *ngrA*-PPTase in biosynthesizing the antimicrobials, we disrupted the *ngrA* gene in the *X. szentirmaii* genome by insertional mutation using a pKNOCK based vector as described in the **Materials and Methods**. After an overlay assay with *Staphylococcus saprophyticus,* the wild type *X. szentirmaii* produced a zone of inhibition. However, the *ΔngrA* mutant strain did not show any anti-microbial properties (**Fig 1C**), further confirming that inactivation of *ngrA* abolishes the synthesis of all NRPS-mediated secondary metabolites (SMs) and bactericidal properties *of X. szentirmaii*.

**Figure 1:**
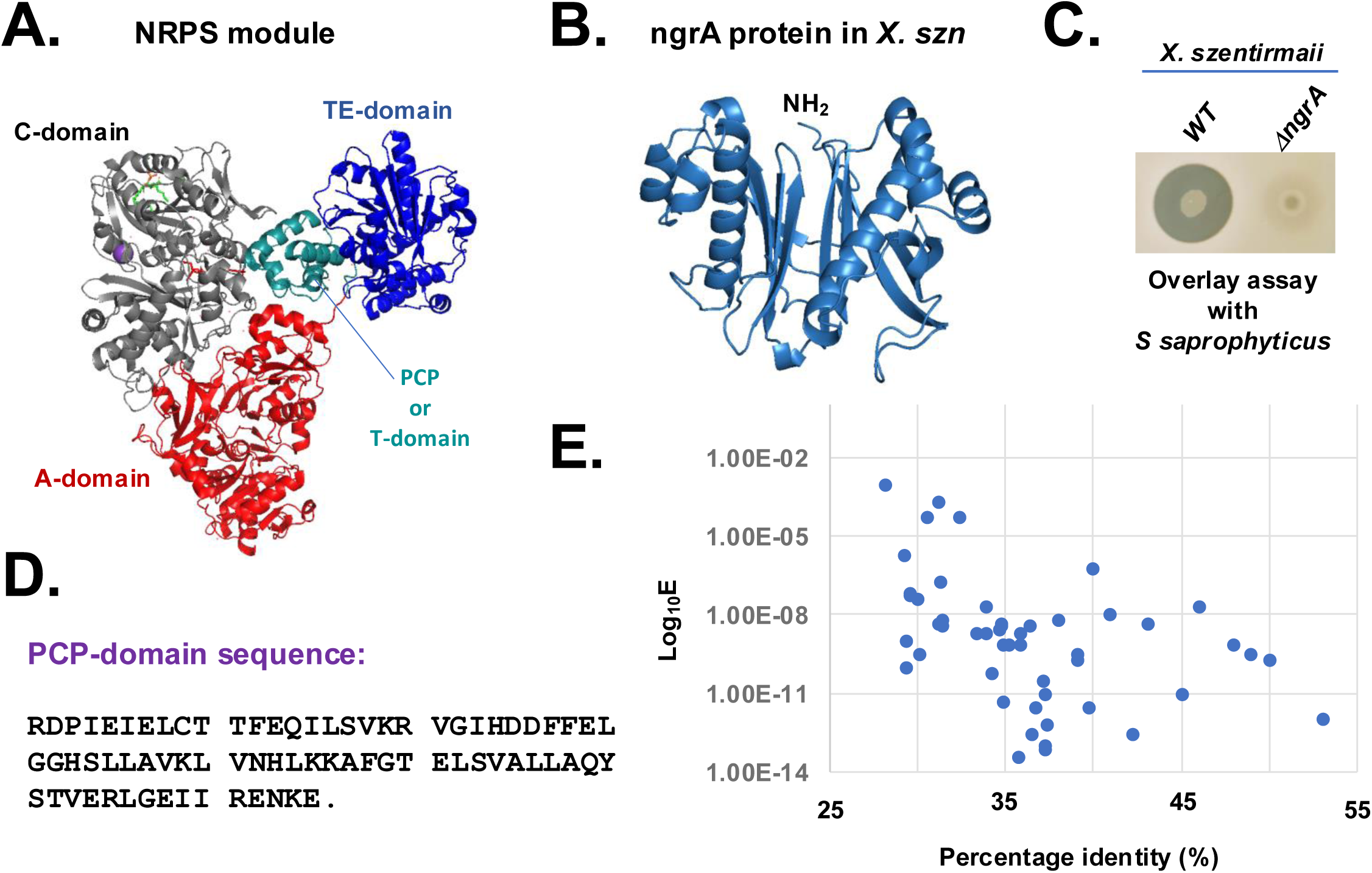
Inactivation of *ngrA* abolishes the synthesis of antimicrobials. (**A**) The crystal structure of *Acinetobacter baumannii NRPS* protein *AB3403 (PDB ID =* 4zxh). The condensation (C)-domain, adenylate (A)-domain, PCP-domain and thioesterase (TE)-domain are shown in grey, orange, red and blue colors, respectively. (**B**) The Alpha-Fold predicted structure of X. szentirmaii (X. szn) ngrA protein. (**C**) The bacterium *X. szentirmaii* (WT or Δ*ngrA* mutant) was spotted on an LB medium and grown overnight. The indicated tester strain *S. saprophyticus* was overlaid and grown at 37°C for 24 hours. The clearance zones indicate the antimicrobial activity. (**D**) Amino acid sequences of the PCP-domain. (**E**) The whole *X. szentirmaii* genome (GenBank accession numbers NIBV01000001 and NIBV01000002) was used to search for genes encoding homologs of the PCP-domain. After the BLAST search, the E-values and percentage identities were retrieved and graphed on a logarithmic scale.

To identify genes encoding protein harboring the PCP domain sequence, the NCBI BLAST search tool was used on the genome of *X. szentirmaii* [21]was analyzed (**Fig 1D** and **Fig 1E**). A total of 64 BLAST-hits were obtained on NRPS, PKS and AMP-binding protein, with a percentage identity >30 and an E-value <10^-4^ (**Fig 1E**). These findings indicated that one or more PCP domains are present in a variety of metabolic enzymes, including ATP-hydrolyzing racemase, dehydrogenase, and non-ribosomal peptide synthetases. We focused on the proteins containing the NRPS-like modules (**Table 1**, protein accession numbers) and considered that two sequences are homologous if they are more than 30% identical over their entire lengths[39]. Then, we analyzed both upstream and downstream sequences of the respective genes to identify their neighboring genes using the SnapGene software (**Table 1**, see GenBank accession numbers). Additionally, the antiSMASH software[40] was used to search for the biosynthetic gene clusters and genes encoding NRPSs in the *X. szentirmaii* genome. Together, our analysis revealed that the PCP domain is distributed among 26 NRPS and PKS genes encoded by 17 operons (referred here to as *ste1* - *ste17,* see **Table1** and **Supplemental Fig 1**).

**Table 1:**
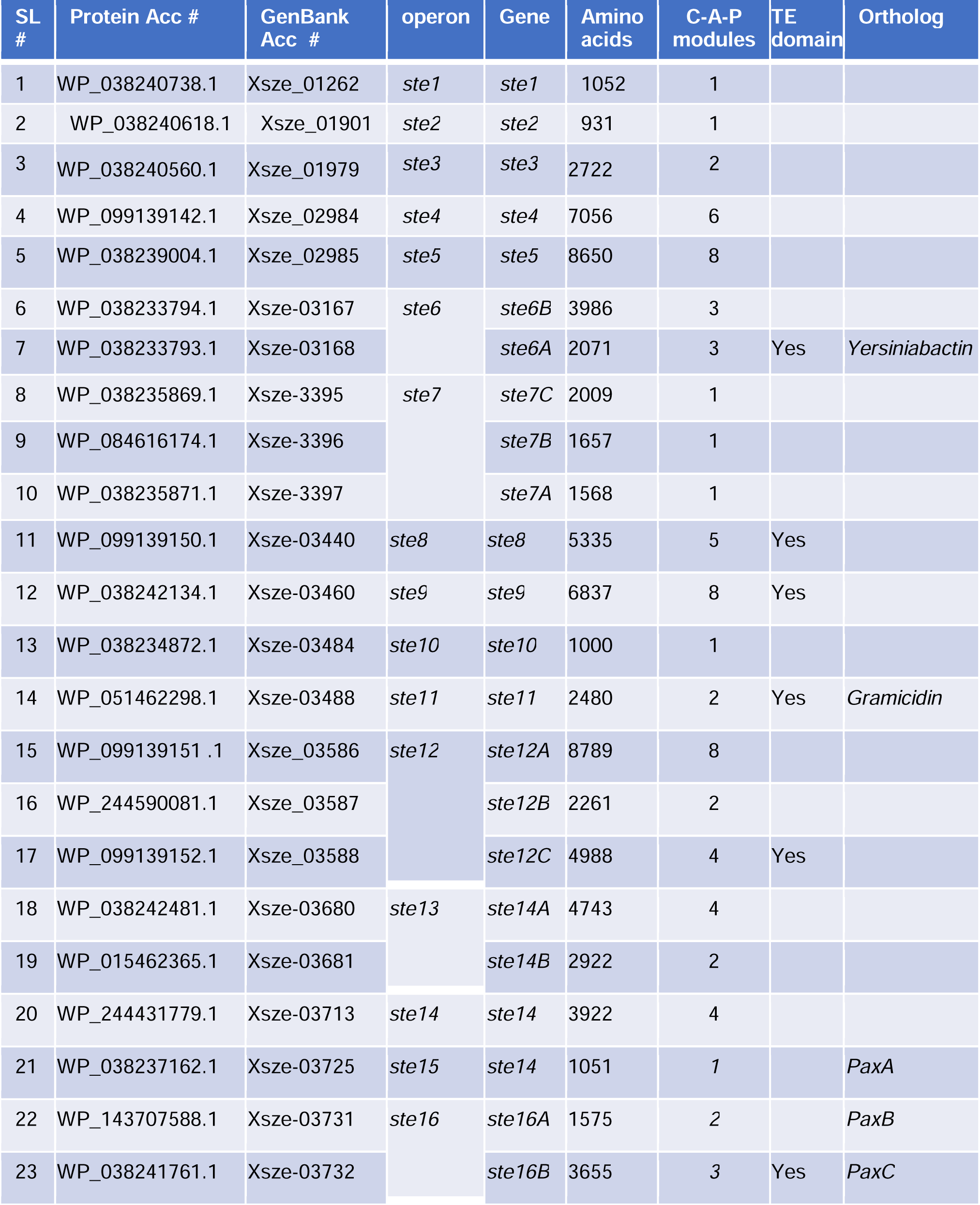
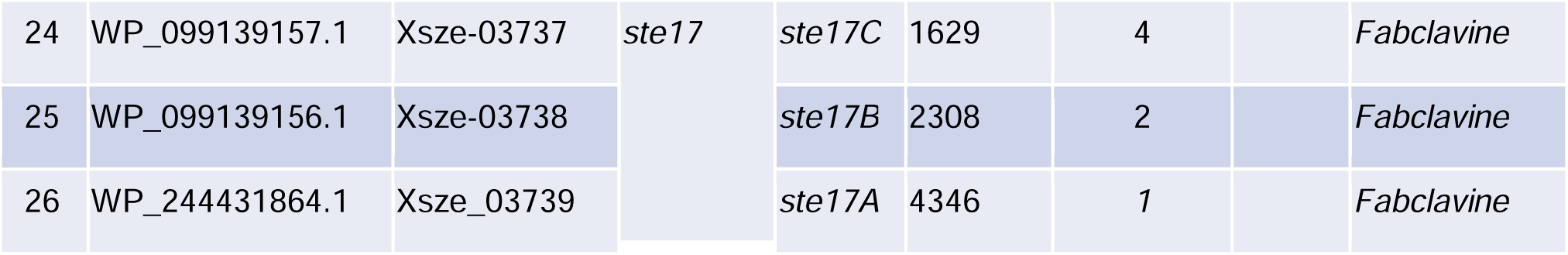
The *X. szentirmaii* genome contains 17 operons that encode NRPS modules.

| SL # | Protein Acc # | GenBank Acc # | operon | Gene | Amino acids | C-A-P modules | TE domain | Ortholog |
| --- | --- | --- | --- | --- | --- | --- | --- | --- |
| 1 | WP_038240738.1 | Xsze_01262 | *ste1* | *ste1* | 1052 | 1 |  |  |
| 2 | WP_038240618.1 | Xsze_01901 | *ste2* | *ste2* | 931 | 1 |  |  |
| 3 | WP_038240560.1 | Xsze_01979 | *ste3* | *ste3* | 2722 | 2 |  |  |
| 4 | WP_099139142.1 | Xsze_02984 | *ste4* | *ste4* | 7056 | 6 |  |  |
| 5 | WP_038239004.1 | Xsze_02985 | *ste5* | *ste5* | 8650 | 8 |  |  |
| 6 | WP_038233794.1 | Xsze-03167 | *ste6* | *ste6B* | 3986 | 3 |  |  |
| 7 | WP_038233793.1 | Xsze-03168 |  | *ste6A* | 2071 | 3 | Yes | *Yersiniabactin* |
| 8 | WP_038235869.1 | Xsze-3395 | *ste7* | *ste7C* | 2009 | 1 |  |  |
| 9 | WP_084616174.1 | Xsze-3396 |  | *ste7B* | 1657 | 1 |  |  |
| 10 | WP_038235871.1 | Xsze-3397 |  | *ste7A* | 1568 | 1 |  |  |
| 11 | WP_099139150.1 | Xsze-03440 | *ste8* | *ste8* | 5335 | 5 | Yes |  |
| 12 | WP_038242134.1 | Xsze-03460 | *ste9* | *ste9* | 6837 | 8 | Yes |  |
| 13 | WP_038234872.1 | Xsze-03484 | *ste10* | *ste10* | 1000 | 1 |  |  |
| 14 | WP_051462298.1 | Xsze-03488 | *ste11* | *ste11* | 2480 | 2 | Yes | *Gramicidin* |
| 15 | WP_099139151.1 | Xsze_03586 | *ste12* | *ste12A* | 8789 | 8 |  |  |
| 16 | WP_244590081.1 | Xsze_03587 |  | *ste12B* | 2261 | 2 |  |  |
| 17 | WP_099139152.1 | Xsze_03588 |  | *ste12C* | 4988 | 4 | Yes |  |
| 18 | WP_038242481.1 | Xsze-03680 | *ste13* | *ste14A* | 4743 | 4 |  |  |
| 19 | WP_015462365.1 | Xsze-03681 |  | *ste14B* | 2922 | 2 |  |  |
| 20 | WP_244431779.1 | Xsze-03713 | *ste14* | *ste14* | 3922 | 4 |  |  |
| 21 | WP_038237162.1 | Xsze-03725 | *ste15* | *ste14* | 1051 | 1 |  | *PaxA* |
| 22 | WP_143707588.1 | Xsze-03731 | *ste16* | *ste16A* | 1575 | 2 |  | *PaxB* |
| 23 | WP_038241761.1 | Xsze-03732 |  | *ste16B* | 3655 | 3 | Yes | *PaxC* |
| 24 | WP_099139157.1 | Xsze-03737 | ste17 | ste17C | 1629 | 4 |  | *Fabclavine* |
| 25 | WP_099139156.1 | Xsze-03738 |  | ste17B | 2308 | 2 |  | *Fabclavine* |
| 26 | WP_244431864.1 | Xsze_03739 |  | ste17A | 4346 | 1 |  | *Fabclavine* |

Based on the sequence homology, five operons (*ste6*, *ste11, ste15, ste16* and *ste17)* likely code for NRPS that catalyze the biosynthesis of orthologs of known compounds, whereas 12 operons (*ste1*, *ste2, ste3, ste4, ste5, ste7, ste8, ste9, ste10, ste12, ste13* and *ste14)* are specific to *X. szentirmaii* with unknown functions. The *ste6* operon is predicted to encode an NRPS that catalyzes the synthesis of a putative *s*iderophore similar to Yersiniabactin[41], whereas the *ste11* operon for the gramicidin-type compound [42]. The *ste15* and *ste16* operons are predicted to encode a composite NRPS that catalyzes the synthesis of antibiotic PAX-peptide [43]. The *ste17* operon appears to contain 11 open reading frames, which is predicted to encode a composite NRPS that catalyzes the synthesis of fabclavine-like compounds (**Table1** and **Supplemental Fig 1** [44]).

Both PKS/NRPS Analysis website (http://nrps.igs.umaryland.edu) and antiSMASH software predict that operons *ste1*, *ste2, ste10* and *15* code for only one modular domain of the NRPS enzyme (**Supplemental Fig 1)**. These observations suggest that these operons alone may not be capable of independently producing functional NRPs and they might work in conjunction with other operon(s) to code a functional NRP. The PKS/NRPS Analysis website predicts that operons ste3, *ste4, ste5, ste8,* and *ste9* respectively code for 2, 6, 8, 5, and 6 C-A-P modular domains (**Table 1**). A terminal thioesterase (TE) domain at the end of these NRPSs suggests that they are likely stand-alone NRPSs. The operons *ste13* and *ste14* encode for 6 and 3 modular domains but lack a TE domain. The *ste13* operon contains 2 NRPS genes (6 modular domains) and the *ste*7 and *ste*12 operons each contain 3 NRPS genes (3 and 14 modular domains, respectively).

A-domain of NRPS modules plays a crucial role in NRP biosynthesis by selecting and activating a specific substrate (typically an amino acid). We utilized two online platforms to predict A-domain substrate(**Table 2**). Some predicted substrates varied between two platforms. For instance, the *ste1*-encoded A-domain was predicted to activate phenylalanine (Phe or F) on the nrpssp.usal.es platform, while the nrps.igs.umaryland.edu platform predicted glycine (Gly or G) as the substrate. These differences highlight the need for further refinement in A-domain substrate prediction. Despite these differences, our analysis revealed that *X. szentirmaii* contains a unique set of 17 NRPS-PKS operons, that are involved in synthesizing a unique repertoire of NRPs, whose functional significance remains to be further characterized.

**Table 2:** Comparison of predicted A-domain substrates from 17 operons.

| Operon | Predicted A-domain substrates |  |
| --- | --- | --- |
|  | https://nrpssp.usal.es | https://nrps.igs.umaryland.edu |
| ste1 | F | G |
| ste2 | F | ? |
| ste3 | F-F | N-W |
| ste4 | T-Y-S-Y-F-T | T-Y-S-?-?-T |
| ste5 | P-N-S-V-F-S-N-G | P-D/N-S-?-?-S-D/N-F |
| ste6 | C <> C | C <> C |
| ste7 | C <> F <> F | ? <> ? <> ? |
| ste8 | V-Y-F-F-Y | V-L-W-W-W |
| ste9 | Y-T-F-V-Y-F | L-T-F-V-?-T |
| ste10 | F | ? |
| ste11 | S-P | ?-? |
| ste12 | P-G-F-V-Y-I-T-V <> V-V <> P-F-P-V | P-G-G-I-L-I-T-V <> V-E <> G-?-P-V |
| ste13 | T-F-T-V <> I-F | T-W-T-E <> I-I |
| ste14 | Y-F-V | V-G-? |
| ste15 | F | G |
| ste16 | V-V <> V-V-V | K-R <> K-K-K |
| ste17 | F-F-N-N <> T-V | ?-?-D/N-D/N <> T-? |

The sign "<>" indicates separation of amino acid encoded by different genes within an operon. The "?" signs indicate unknown amino acids.

**Table 3:**
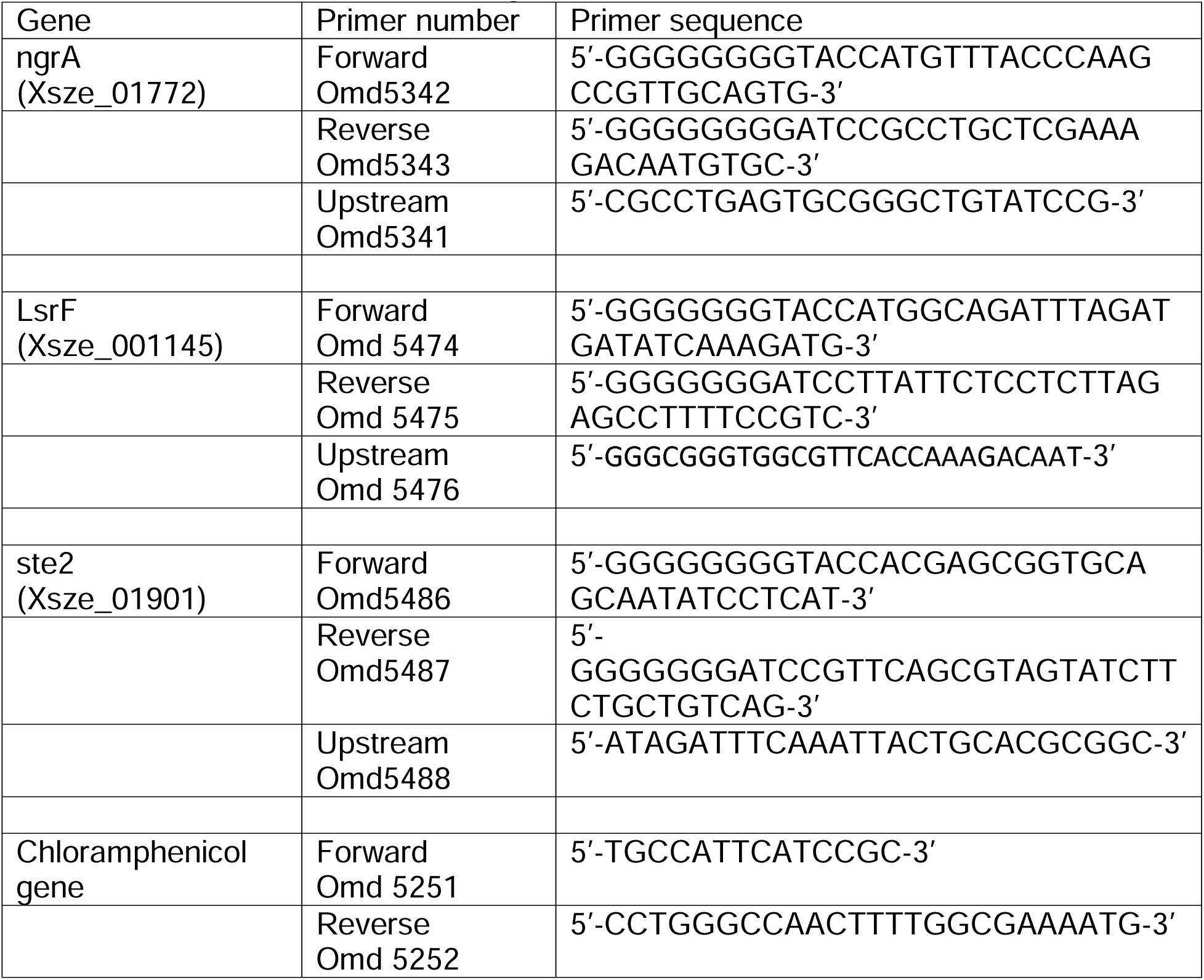
Primers used in this study.

### 2. *X. szentirmaii* and *X*. *nematophila* produces antimicrobials against pathogenic bacteria

*Xenorhabdus nematophila* has been studied extensively for its ability to secrete antimicrobial compounds. Here, we sought to compare the differences in antimicrobial secretion between *X. nematophila* and *X. szentirmaii* using an overlay assay with the tester strains *S. saprophyticus* and *E. coli* as described in the **Materials and Methods**. A large zone of inhibition was observed surrounding the *X. szentirmaii* when overlaid with *S. saprophyticus* (**Fig 2A**), and even more prominent inhibition zone was seen when overlaid with *E. coli* when compared to *X. nematophila* (**Fig 2A**). These results further confirmed that *X. szentirmaii* produces a higher level of antimicrobials [16].

**Figure 2:**
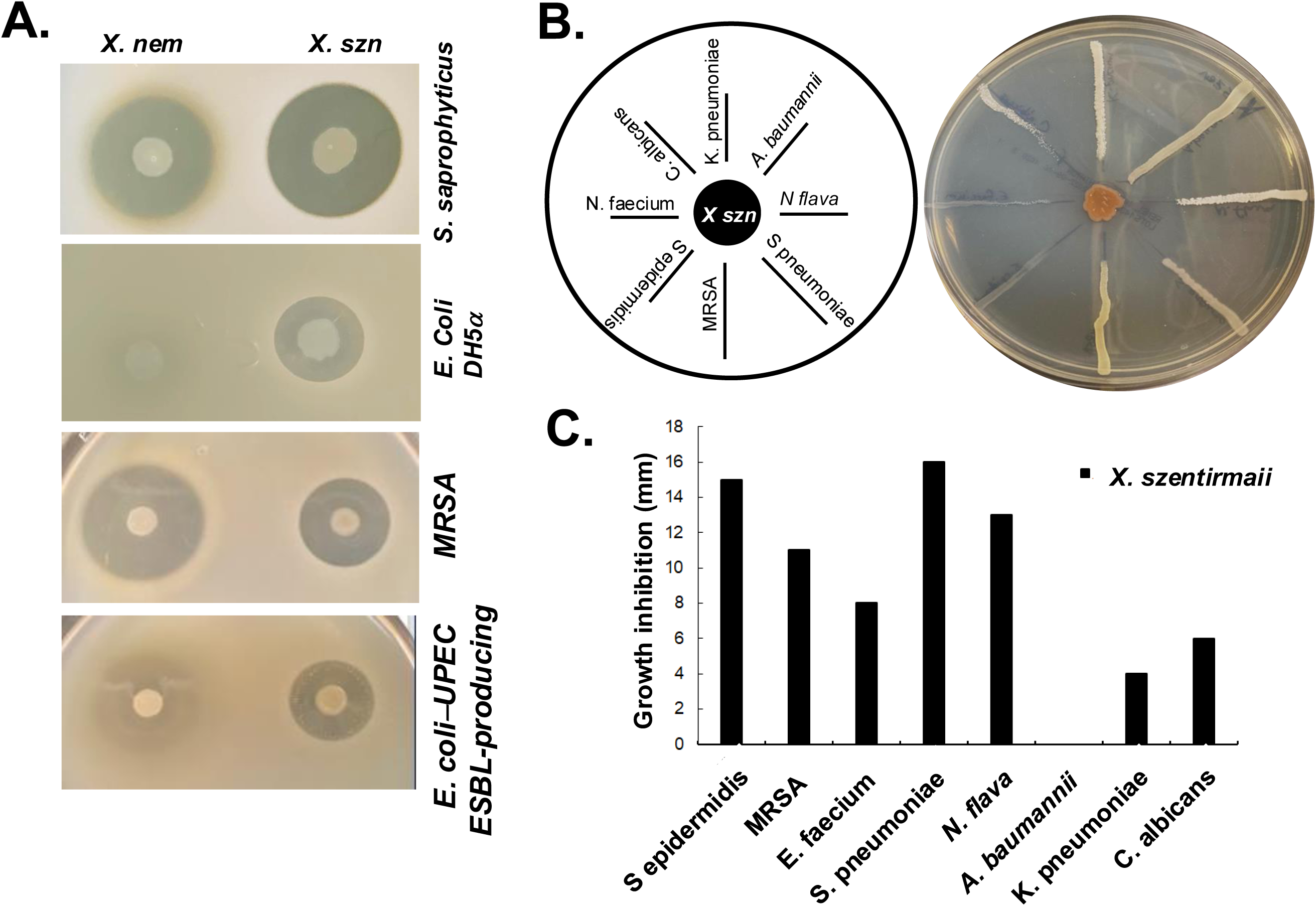
X nematophila and X. szentirmaii are natural resources of antibiotics. (**A**) The bacteria *X. nematophila* and *X. szentirmaii* were subjected to overlay assays with four tester strains: *S. saprophyticus, E. coli,* MRSA and a clinical ESBL-producing UPEC strain. (**B**) The bacterium *X. szentirmaii* was grown for 48h before cross-streaking with the indicated microorganisms. The distance of inhibition was measured and recorded in a bar diagram (**C**).

To assess the antimicrobial effects against antimicrobial resistant organisms recognized as “serious threats” by the CDC, the methicillin-resistant *Staphylococcus aureus* (MRSA) and an extended-spectrum beta-lactamase (ESBL)-producing uropathogenic *E. coli* (UPEC) (**Fig. 2A**) were tested. Both *X. szentirmaii* and *X. nematophila* demonstrated strong anti-microbial effects against MRSA and ESBL-producing *UPEC*, with *X. szentirmaii* exhibiting stronger effects against the latter compared to *X. nematophila* (**Fig. 2A**).

To further determine if *X. szentirmaii secretes* antimicrobial compounds capable of inhibiting growth of a variety of ESKAPE (*<u>E</u>nterococcus*, *<u>S</u>taphylococcus*, *Klebsiella, <u>A</u>cinetobacter, <u>P</u>seudomonas*, *<u>E</u>nterobacter*) pathogens commonly associated with resistance, we conducted a cross-streak assay (**Fig 2B**). *X. szentirmaii* was grown for 48h on an LB plate, followed by streaking seven pathogenic bacteria: (*Klebsiella pneumoniae, Acinetobacter baumannii, Neisseria flava, Streptococcus pneumoniae, methicillin-resistant Staphylococcus aureus, Staphylococcus epidermidis, Enterococcus faecium)* and a clinically relevant fungus *(Candida albicans),* as illustrated in **Fig 2B**. The plate was then grown at 37°C for 48 hours. Except for the pathogen *Acinetobacter baumannii*, various zones of inhibition were evident for all tested bacteria and the yeast strain (**Fig 2B and C**), supporting potential application of *Xenorhabdus* bacteria against AMR pathogens.

### 3. *X. szentirmaii* produces antimicrobials during the stationary phase

To determine the specific growth stage of antibiotic production by *X. szentirmaii*, cell-free supernatants (CFSs) were collected from the log- and stationary-phases of bacterial cultures (**Fig 3A**) and subjected to overlay assays with the tester strain *S. saprophyticus*. Zones of inhibition were observed surrounding the CFSs collected at the stationary phase of cultures compared to log phases (**Fig 3B**). These data indicated two possibilities: (I) *X. szentirmaii* did not secrete antimicrobial compounds during the log phase and/or (II) number of cells at the log phase were too low to produce antimicrobials.

**Figure 3:**
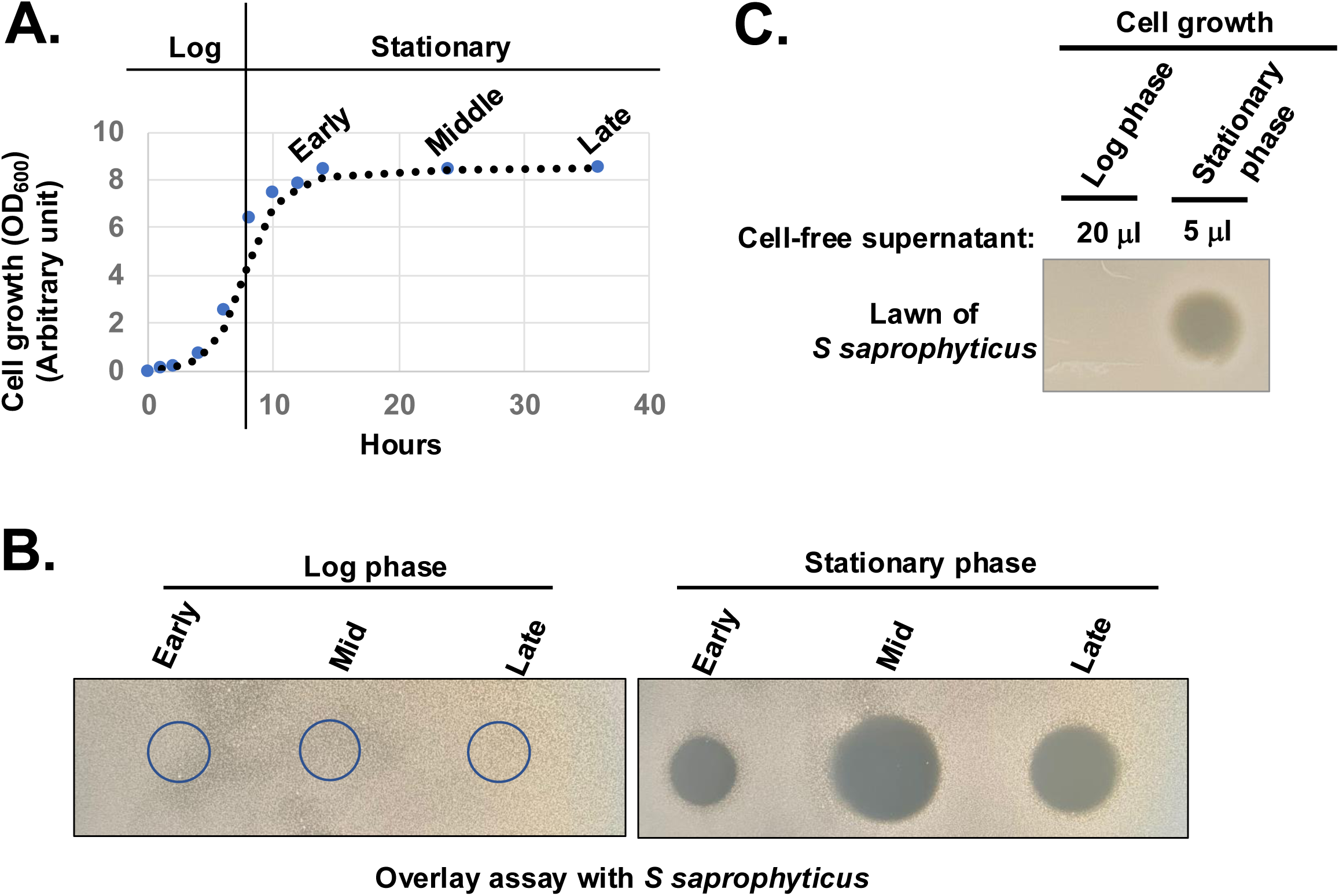
*X. Szentirmaii* produces antimicrobials in the stationary phase. (**A**) The bacterium *X. szentirmaii* was grown for the indicated time and the optical density (OD) was measured at 600 mm. The experiments were repeated at least three times. One representative data are shown. (**B**) Cell-free supernatants were collected from both log and stationary phases, filter-sterilized, spotted (5μl) on an agar plate, and subjected to overlay assay with the bacterium *S. saprophyticus*. The clearance zone indicates antibacterial activity. (**C**) The 20μl or 5μl of cell-free supernatants containing equal number of cells were collected, concentrated, and spotted on the LB medium and subjected to overlay assay.

To test the above possibilities, we collected CFSs containing an equal number of cells from both log and stationary phases, considering the lower cell density at the log phase (OD_600_=∼0.5) compared to the stationary phase (OD_600_ <1.5). CFS were then dried by speed vacuum, resuspended in equal volume of LB medium and spotted on the LB medium for the overlay assay with the tester strain *S. saprophyticus*. Zones of inhibition were observed surrounding the CFSs collected only at the stationary phase of cultures (**Fig 3C**). Taken together, these observations suggest that a certain cell density is required for the secretion of antimicrobial compounds and the secretion may be regulated by certain components of quorum sensing regulators.

### 4. Genome-wide analysis of transcripts expressed during log and stationary phases

In *X. szentirmaii*, the biosynthetic genes of secondary metabolites appear to be dominated during the late log and stationary phases (**Fig 3**). To understand the molecular details of the biosynthesis, secretion, and/or mode of action of antimicrobials, we performed a comparative analysis of transcripts expressed during log and stationary phases of growth to identify the deferentially expressed genes. Total RNA was isolated from *X. szentirmaii* grown at log and stationary phases (**Fig 4A**, four biological replicates referred here Dey1, Dey2, Dey3 and Dey4 for the log phase and Dey5, Dey6, Dey7 and Dey8 for the stationary phase, **Supplemental Table 1**). The equal amount of total RNA was subjected to RNA sequencing analysis. The RNA-seq data was then used to generate the PCA (principal component analysis) plot, which displayed distinct clustering (**Fig 4B**), indicating that there are discernable differences in gene expression between the log and stationary phases, and that the biological conditions associated with each phase lead to distinct gene expression pattern.

**Figure 4:**
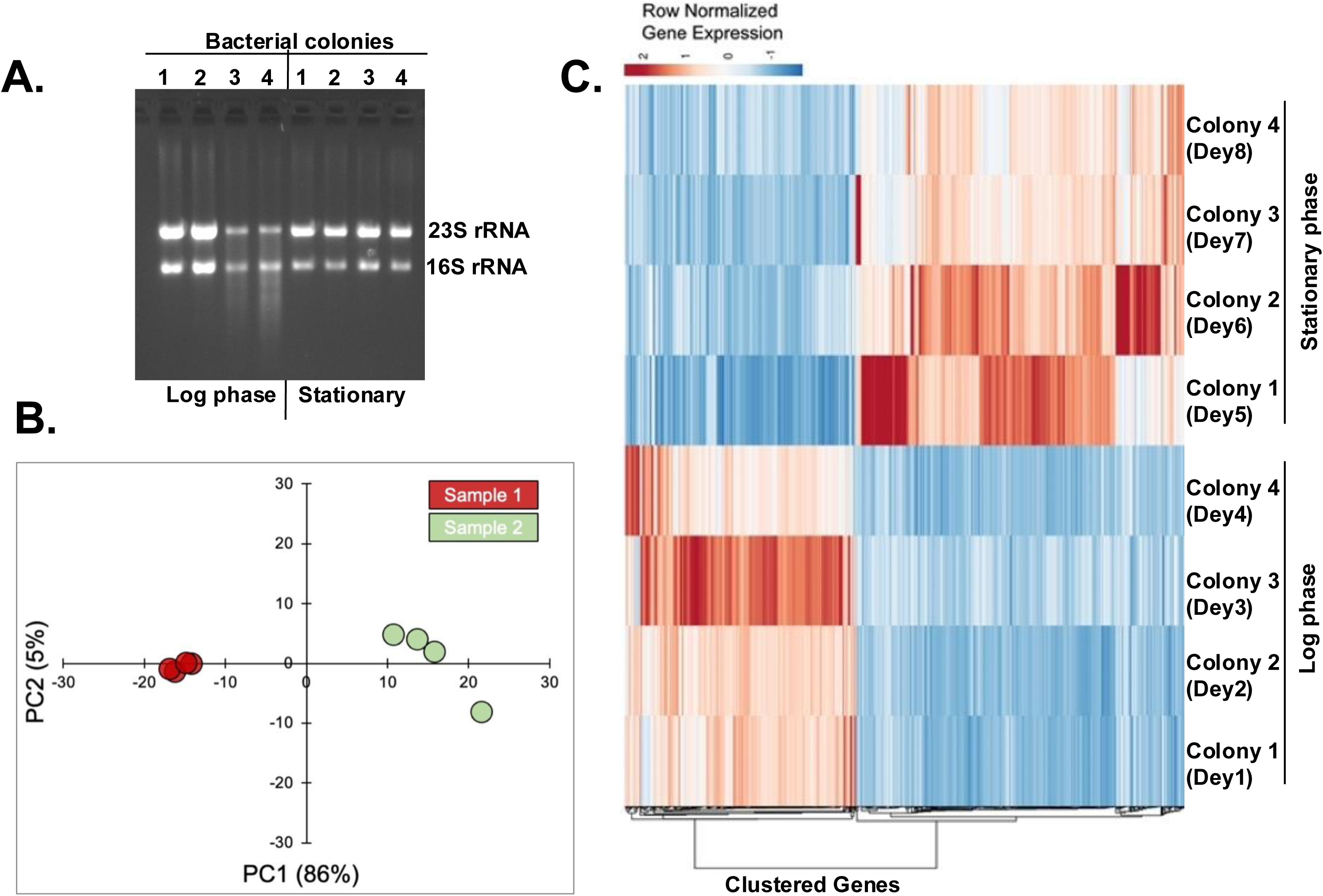
Analysis of transcripts expressed during log and stationary phases. **(A)** Four bacterial colonies were grown till the log and stationary phases. Total RNAs were isolated and run in an agarose gel. **(B)** The principal component analysis (PCA) plot shows two distinct clusters of samples from log-phase (green dots) and stationary-phase (red dots). **(C)** Heatmap of differentially expressed genes. The heatmap shows the row normalized TPM (Transcripts per million) values of 637 differential genes with FDR < 0.05 and absolute log_2_FC above >1.

The draft genome sequence featuring 3810 genes of *X. szentirmaii* DSM 16338 was leveraged as a reference[21]. Our RNA-seq analyses identified 3446 gene products. Approximately 637 genes were upregulated during the log or stationary phase (absolute log_2_ fold-change more than1) (**Fig 4C, Supplemental Table 2**). Notably, As shown in **Fig 5A**, the functions of 118 genes remained uncharacterized. The remaining gene products include transcription factors (e.g., LysR,), synthetases (e.g., dethiobiotin synthetase), transferases (e.g., GNAT family acetyltransferase), metabolic kinases (e.g., phosphoenolpyruvate carboxy-kinase), phosphatases (e.g., acid phosphatase and phosphotransferase), reductases (e.g., oxidoreductase), transporters (e.g., putative metal cation transporter P-type ATPase CtpV, permeases (e.g., ABC transporter permease), lyases (e.g., aspartate ammonia-lyase), chaperones (e.g., ClpB) and many membrane proteins (e.g., tricarboxylic transport membrane protein). Notably, no significant increase in transcripts encoding replication factors was observed. These findings suggest that these genes are involved in regulating various cellular processes, including transcription, transport, metabolisms, and biosynthesis of secondary metabolites and their modifiers. Together, these findings indicate that *X. szentirmaii* reprograms its physiological state upon reaching a certain cell density.

**Figure 5:**
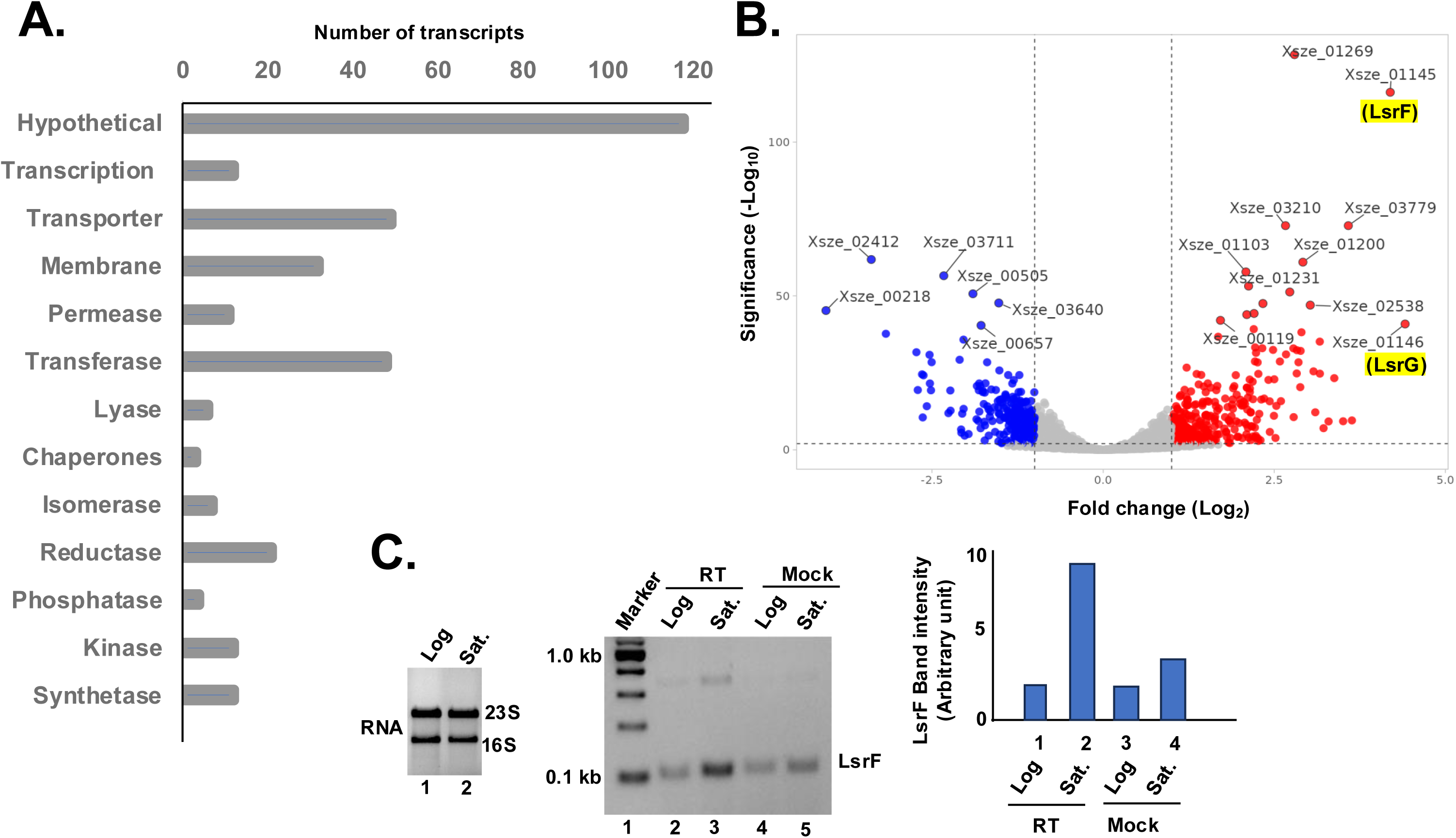
Upregulation of quorum sensing activities during the stationary phase. **A.** The bar diagram shows the number of up-regulating genes during the stationary phase are classified based on their physiological functions. **B.** Total RNA was isolated from the log and stationary phases of *X. szentirmaii* and subjected to RNA-seq analysis. Up and down regulated genes are shown in red and blue color, respectively, in the volcano plot. **C.** (Left panel) Total RNA was isolated from the log and stationary phases of *X. szentirmaii* and run on an agarose gel. (Middle panel) cDNA was synthesized by reverse transcriptase (RT) from total RNA isolated from log and saturated (Sat.) phases and subjected to RT-PCR with the LsrF-specific primers. The PCR products were run in an agarose gel. A control reaction without RT (Mock) was also carried out. (Right panel) The band intensities of the PCR products were measured NIH-image J software and shown in a bar diagram.

The top 10 upregulated genes during the stationary phase (**Fig. 5B**) are *Xsze-01146 (*encoding autoinducer-2 degrading protein **LsrG**), *Xsze-01145 (autoinducer-2 aldolase **LsrF**)*, *Xsze-01251* (fumarate reductase subunit D), *Xsze-03779* (urocanate hydratase), *Xsze-03299* (dethiobiotin synthetase), *Xsze-03232* (putative alcohol-acetaldehyde dehydrogenase), *Xsze-01253* (fumarate reductase), *Xsze-01252* (fumarate reductase subunit C), *Xsze-03764* (tricarboxylic transport membrane protein) and *Xsze-01254* (fumarate reductase flavoprotein subunit). Further genome analysis showed that *lsrF* and *lsrG* are two conserved structural genes of the *lsr* operon in *X. szentirmaii*, which is likely regulated by the transcription factor LsrR (**Supplemental Figure 2**). However, the functional significance of *lsr* operon in *X. szentirmaii* is yet to be determined.

To confirm that the *lsrF* gene expression was increased during the stationary phase, we conducted the reverse transcriptase (RT)-PCR analysis. Total RNA was prepared from both the log phase and the stationary phase of cell growth (**Fig 5C**) and subjected to RT-PCR analysis by two gene-specific primers of *lsrF*. The increased amount of PCR-amplified product (5-fold) in the presence of RT (**Fig 5C**, middle panel, lane 3, compared with the lane 5 and the right panel) confirmed that the expression of *lsrF* gene is induced during the stationary phase. These data indicate that LsrF and LsrG may have a distinct function in regulating antibiotic production by *X. szentirmaii*.

Consistent with the previous studies[45], our genome analysis further revealed that *lsrF* and *lsrG* are the only components of the *lsr* operon involved in regulating the QS system in *X. szentirmaii*. (**Supplemental Figure 2**). QS-mediated gene regulation arises from regulatory networks built upon the core mechanisms of AI production and detection. Thus, it appears that specific patterns may become apparent when investigating the role of QS network in antimicrobial production. It is plausible that certain regulatory systems within the QS network may have evolved to specifically control NPR biosynthesis. Future molecular studies will focus on the role of LsrF in NPR biosynthesis.

### 5. Increased expressions of ste2, ste10, ste14, ste17 and *ngrA* genes during the stationary phase

*X. szentirmaii* produced antimicrobials during the saturated phase when cells reached a certain cell density (**Fig 3**), and this production of antimicrobials depended on the function of NgrA-PPTase (**Fig 1**). In this study, we also identified that at least 17 NRPS operons regulated the expression of antimicrobial NRP (**Table 1**). However, the RNA-seq data showed that only the expression of *ste10* (*Xsze-03484*) gene was increased (log_2_fold-change = 1.30 with a p-value 4.91E-08, **Supplemental Table 2**). These results suggested that the RNA-seq experiment may have failed to fully detect these NRPS transcripts likely due to their large size and/or short half-life. Additionally, it was possible that a 2-fold increase in transcripts might have not been captured by the high-throughput analysis. Therefore, we individually examined the raw FPKM (<u>F</u>ragments <u>P</u>er <u>K</u>ilobase of transcript per <u>M</u>illion mapped reads) counts of each *ste* operon (**Supplemental Figure 1**) and analyzed their expression levels (**Fig 6**).

**Figure 6:**
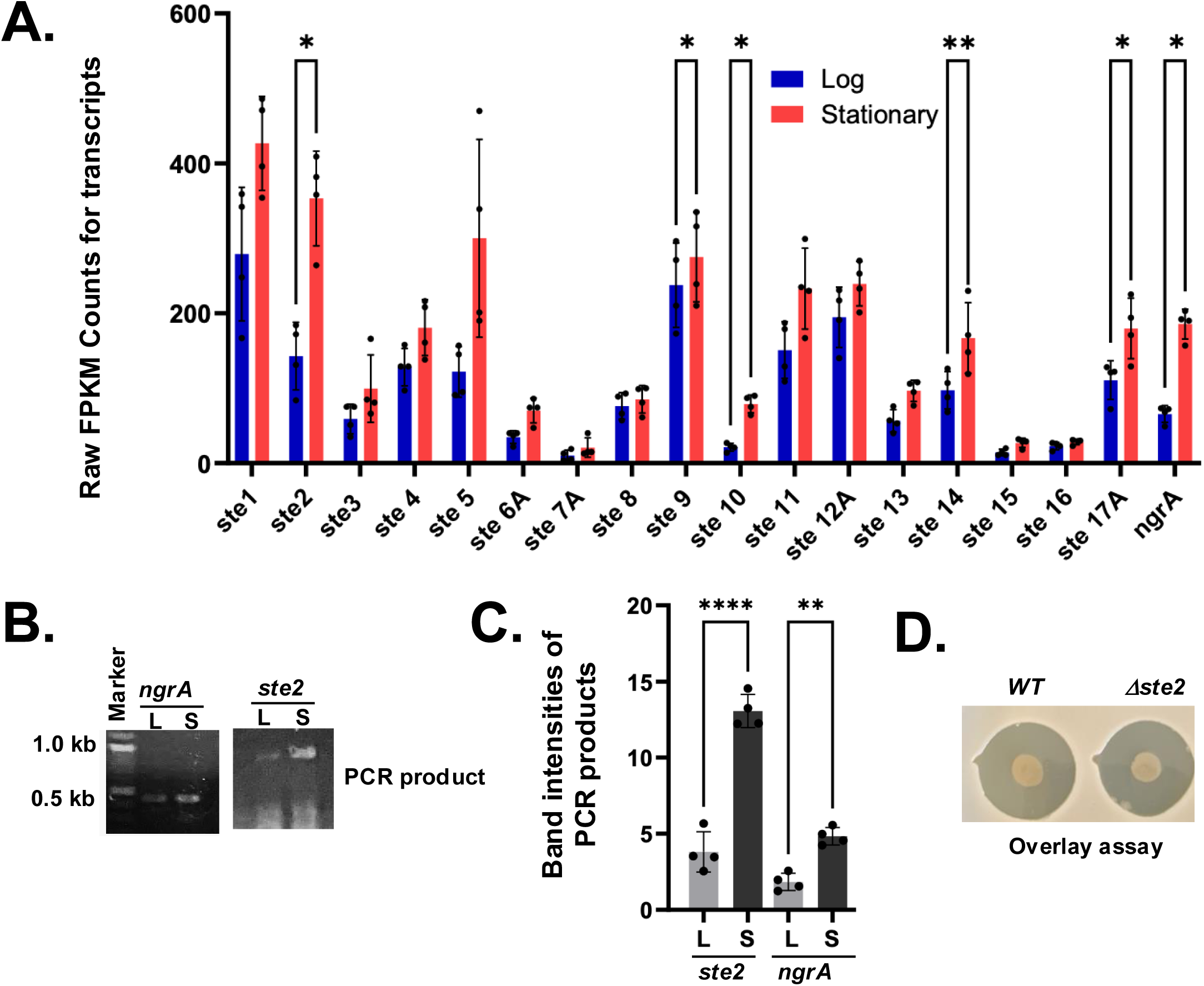
Upregulation of operons ste2, ste9, ste10, ste14A and ste17A during the stationary phase. **A.** The average FPKM counts of the indicated *ste* genes are shown in a bar diagram with standard error (**p-value<0.001, paired t-test). **B.** cDNA was synthesized by reverse transcriptase (RT) from total RNA isolated from log (L) and saturated (S) phases and subjected to RT-PCR with primers designed from the indicated ngrA and *ste2* operons. **C.** The PCR-amplified DNA bands were measured by ImageJ software and indicated in a bar diagram with standard error (*p-value<0.001, paired t-test). **D.** Wild type (WT) and *Δste2* strains of *X. szentirmaii* were subjected to overlay with the tester strain *S. saprophyticus*. The experiments were repeated at least three times. Results from one representative experiment data are shown.

Our analysis showed that there were significant variations in the FPKM counts for *ste* transcript levels. The average FPKM counts at log and stationary phases are 295 and 412 for *ste1*, 151 and 383 for *ste2*, 63 and 116 for *ste3*, 143 and 184 for *ste4*, 126 and 934 for *ste5*, 40 and 66 for *ste6*, 12 and 26 for *ste7*, 12 and 26 for *ste8*, 77 and 90 for *ste8*, 253 and 287 for *ste9*, 24 and 81 for *ste10*, 154 and 264 for *ste11*, 224 and 251 for *ste12A*, 64 and 107 for *ste13*, 107 and 189 for *ste14*, 15 and 30 for *ste15*, 24 and 26 for *ste16A*, and 124 and 198 for *ste17A* (**Supplemental Table 1**). These numbers show an overall increase in each NRPS transcript at the stationary phase. However, significant values were obtained for *ste2* (∼2.5-fold with a p-value = 0.029, paired t-test), ste9 (∼0.85-fold with a p-value = 0.039, paired t-test), *ste10* (3-fold with a p-value = 0.019, paired t-test), ste14A (∼2-fold with a p-value =0.005, paired t-test) and ste17A (1.5-fold with a p-value = 0.037, paired t-test) (**Fig 6A**). Although the ste5 transcript level was increased by ∼7-fold, a higher p-value (0.1962, paired t-test) indicated that this increased expression was not statistically significant (not analyzed). The increase in the NRPS transcript levels was consistent with the increase in the *ngrA* transcript level (*Xsze_01772*), which showed an average increase of ∼3.5-fold with a p-value of 0.03 (paired t-test). Together, these findings suggest that the regulatory function and turn-over of the NRPSs might vary considerably based on their length and amino acid composition.

To validate the RNA-seq data, we performed reverse-transcript (RT)-PCR analysis targeting the operons ngrA and *ste2.* cDNA was synthesized from the total RNA isolated from both log and stationary phases and amplified using gene-specific primers. Consistent with the RNA-seq data, we found that transcription of ngrA and *ste2* was significantly increased in bacteria grown at their stationary phase (**Fig 6B** and **Fig 6C**). Taken together, our results suggest that the expressions of ngrA and *ste2* increased during stationary phase, coinciding with the antibiotic production.

In this study, to further investigate the role of *ste operons* in antimicrobial secretion, we specifically disrupted the *ste2* operon by insertional mutagenesis using a pKNOCK based vector (**Materials and Methods**), generating a *Δste2* strain. An overlay assay with *Staphylococcus saprophyticus* was then conducted to compare the *Δste2* strain and the wild type of strain. No difference in the zone of inhibition was observed between WT and *Δste2* strains (**Fig 6D**). These results indicate that all *ste* operons likely function collectively to regulate antimicrobial secretion. Future studies will focus on disrupting each *ste* operon individually and in combination to evaluate their relative contribution to antimicrobial secretion.

## Discussion

We focused our research on *X. szentirmaii,* a species that grows faster than its close relatives and produces more potent antimicrobial compounds against AMR (**Fig 2**). Genome analysis of the *X. szentirmaii* genome identified 17 operons (**Table 1**), encoding NRPS and PKS enzymes (*ste1*-*ste17)*. RNA-seq analysis revealed that the expression of these operons increased during the stationary phase of its growth when *X. szentirmaii* secretes antibiotics. Furthermore, RNA-seq analysis identified that the quorum sensing (QS)-regulator LsrF is significantly elevated during the stationary phase and a key factor in regulating the antibiotic production (**Fig 5**).

The *X. szentirmaii* genome contains at least 24 NRPS and/or PKS genes, which are distributed among 17 operons (**Table 1**). Based on sequence homology, some operons likely code for orthologs of known compounds. For example, *ste6* operon is predicted to encode NRPSs to biosynthesize a *s*iderophore similar to Yersiniabactin[41], while the operon *ste11* to biosynthesize the gramicidin-type compound [42]. Both *ste15* and *ste16* operons are predicted to encode enzymes for synthesis the antibiotics PAX-peptide [43]. The operon 17 is predicted to encode NRPSs to biosynthesize Fabclavine-like compound. The fabclavine was first discovered in *X. budapestensis* and *X. szentirmaii* as a hexapeptide-*polyketide*, with a corresponding 50kb biosynthesis gene cluster consisting of non-ribosomal peptide synthetases and one polyketide synthase [44]. Later, several isoforms of fabclavine and their corresponding biosynthesis gene clusters were reported, which have generated significant attention due to their broad-spectrum bioactivity against Gram-positive and -negative bacteria, fungi, and protozoa[44, 46, 47]. Recently it has been shown that Fabclavine from *X. szentirmaii* can kill the larva of the mosquito Aedes albopictus [48]. Here, we report that the fabclavine-like compound in *X. szentirmaii* is likely to be a heptapeptide, with a corresponding 28.6 kb biosynthesis gene cluster consisting of five genes (**Supplemental Figure 1**).

The products and functions of the rest of the operons are currently unknown. Some operons encode only one modular domain of the NRPS enzyme, indicating that these operons alone may not be capable of independently producing functional NRPs, but instead collaborate with other operon(s) to synthesize complete and functional NRPs. Additionally, some operons lack a terminal TE domain, indicating that they may work in conjunction with other genes that contains a TE domain to produce an NRP. It remains to be determined whether the five compounds reported in *X. szentirmaii* (i.e., Xenematide, GameX peptide, Rhabdopeptide, Xenoamicin, Szentiamide) [21] are synthesized by a single operon or by NRPS encoded across multiple *ste* operons.

It appears that there are five stand-alone operons for NRPSs in *X. szentirmaii*. These operons are *ste4* (encoding an NRP of six amino acids)*, ste6* (encoding an NRP of hexa-peptide), *ste8* (a pentapeptide NRP), *ste9* (another hexa-peptide NRP) and ste12 (encoding a fourteen-peptide NRP, **Supplemental Figure 1**). Each of these operons likely encodes a complete NRPS, making them attractive candidates for engineering and producing the natural bioactive NRP antibiotics, either within *X. szentirmaii* or in a heterologous *E coli* strain [49], to combat the growing issue of antibiotic resistance.

Taken together, the *X. szentirmaii* genome contains a unique set of NRPSs that potentially synthesize a unique diversity of antimicrobial non-ribosomal peptides. However, the functional characterization of these operons at the molecular levels remains to be determined. Notably, the disruption of the *ste2* operon did not impair the ability of *X. szentirmaii* to inhibit the *S. saprophyticus* growth (**Fig 6D**), suggesting that all *ste* operons might function collectively in mediating both intra- and interspecies competition. Ongoing research aims at disrupting each *ste* operon individually and in combination to evaluate their role in producing antimicrobial compounds. Additionally, future studies will aim to strategically express and engineer these NRPS operons in *E coli*.

Another key finding in our research is that antibiotic secretion in *X. szentirmaii* is regulated by QS components. The LuxR/LuxI-type QS system, which facilitates cell-cell communication and bioluminescence in more than 100 Gram negative bacteria[50], has been shown to regulate the biosynthesis of the non-ribosomal lipopeptide keanumycin D in *Pseudomonas nunensis* [51]. Additionally, it is reported that the *NRPS* gene regulates the synthesis of some QS signal molecules in *S. odorifera* [52]. In *X. szentirmaii*, the QS consists of only LsrF and LsrG (**Supplemental Figure 2**). Comparative analysis of transcripts expressed in cells at exponential and stationary growth phases showed a significant increase in the expression of *lsrF* and *lsrG* genes during the stationary phase, coinciding with the antibiotic production (**Fig 5**). These findings suggest that the LsrF plays a crucial role in antibiotic secretion. In line with the previous reports [51, 52], our results indicate that the QS system provide a substantial advantage to bacterial communities producing non-ribosomal peptides. This study provides new insights into microbiome research, contributing to the development of novel antibiotics.

The *lsr* operon in both *E coli* and *Salmonella enterica* contains multiple genes encoding the components for the uptake and manipulation of auto-inducers (AI)-2[53]. Specifically, within this operon, genes *lsrA*, *lsrB*, *lsrC*, and *lsrD* encode proteins involved in the transport of AI-2, while *lsrF*[54] and *lsrG*[55] encode proteins involved in the processing of the phosphorylated AI-2, thereby modulating quorum-sensing-regulated bacterial behaviors. The *lsr* operon is further regulated by its adjacent *lsrR* and *lsrK* genes, encoding a repressor and a metabolic kinase[56, 57]. Unlike *E coli,* the *lsr* operon in *X. szentirmaii* is distinct, it contains only *lsrF* and *lsrG* genes, lacking LsrB, a known receptor for AI-2[58]. Consistently, it has been reported that AI-2 production is not involved in the global regulation of natural product biosynthesis in *Xenorhabdus*[45]. However, we found that there is a 90% similarity between LsrF proteins in *E. coli* and *X. szentirmaii,* implying that these proteins have a common function in both bacteria (**Supplemental Figure 2**). The crystal structure shows that LsrF encodes a protein of ∼32 kDa that folds into a homo-decameric enzyme and catalyzes the formation of key metabolites dihydroxy acetone phosphate and acetyl CoA[54] (**Supplemental Figure 2**). However, the roles of these metabolites in the gene regulation or QS are not clear.

Our data indicate that the QS-regulators LsrF and LsrG may have a distinct function in regulating the antibiotics produced by *X. szentirmaii*. QS is an important mechanism to regulate the expression of secondary metabolites, ensuring that these compounds are produced when there is a sufficient population of microorganisms. Specifically, pathogenetic bacteria frequently rely on QS to synchronize the expression of virulence factors, leading to biofilm formation and invasion[59]. Thus, a deeper understanding of LsrF and LsrG is needed to further understand their roles in quorum sensing and antibiotic secretion, which can be applied to other pathogenic bacteria.

## Conclusions

AMR is a major health crisis with growing numbers of bacteria and fungi becoming resistant to existing antibiotics [1]. It is estimated that AMR costs up to $40 billion in direct costs alone in the United States[60], which is primarily driven by some of the following WHO priority pathogens: *Enterococcus faecium, Staphylococcus aureus, Klebsiella pneumoniae, Acinetobacter baumannii*, *Pseudomonas aeruginosa and Escherichia coli* [61]. Continuous efforts have been made to identify new antibiotics from natural resources, including the well-studied bacterial genus *Streptomyces* [4]. However, a promising yet relatively untapped natural resource of antibiotics lies within the Gram-negative *Xenorhabdus* bacteria, which secretes numerous antimicrobial compounds [5]. None of those compounds have been commercial utilized due to several technical challenges and a limited understanding of bacterial molecular biology. This highlights the need for fundamental molecular studies on these bacteria. In this study, we demonstrate that *X. szentirmaii* produces several NRPSs and PKSs through 17 operons and quorum sensing regulators. These NRPSs and PKSs biosynthesize potent antimicrobial NRPs, which are unique and substantiate the potential of *Xenorhabdus szentirmaii* as a promising natural source for future antimicrobial discoveries in a society that needs new antibiotics to combat AMR.

## Acknowledgements

DOV was supported by a fellowship from the Office of Undergraduate Research (OUR), UW-Milwaukee. The RNA sequencing experiments were supported by the Great Lakes Genomics Center at the University of Wisconsin Milwaukee (RRID:SCR_017838) with Illumina library preparation and sequencing.

## Author contributions

RD conceptualized, performed experiments, and wrote the paper, DOV performed the RT-PCR analysis, AC performed bioinformatics analysis, KM and JU performed experiments, SM conceptualized the experiments and wrote the paper, SF conceptualized and wrote the paper, TS conceptualized, performed the experiments, and wrote the paper, MD conceptualized, performed experiments, and wrote the paper. All authors edited the paper.

## Notes

### Competing Interest Statement

The authors have declared no competing interest.

## References

1. Brown, E.D. and G.D. Wright, Antibacterial drug discovery in the resistance era. Nature, 2016. 529(7586): p. 336–43.

2. Antimicrobial Resistance, C., Global burden of bacterial antimicrobial resistance in 2019: a systematic analysis. Lancet, 2022. 399(10325): p. 629–655.

3. Reygaert, W.C., An overview of the antimicrobial resistance mechanisms of bacteria. AIMS Microbiol, 2018. 4(3): p. 482–501.

4. Chevrette, M.G., et al., The antimicrobial potential of Streptomyces from insect microbiomes. Nat Commun, 2019. 10(1): p. 516.

5. Forst, S., et al., Xenorhabdus and Photorhabdus spp.: bugs that kill bugs. Annu Rev Microbiol, 1997. 51: p. 47–72.

6. Herbert, E.E. and H. Goodrich-Blair, Friend and foe: the two faces of Xenorhabdus nematophila. Nat Rev Microbiol, 2007. 5(8): p. 634–46.

7. Booysen, E. and L.M.T. Dicks, Does the Future of Antibiotics Lie in Secondary Metabolites Produced by Xenorhabdus spp.? A Review. Probiotics Antimicrob Proteins, 2020. 12(4): p. 1310–1320.

8. Snyder, H., et al., New insights into the colonization and release processes of Xenorhabdus nematophila and the morphology and ultrastructure of the bacterial receptacle of its nematode host, Steinernema carpocapsae. Appl Environ Microbiol, 2007. 73(16): p. 5338–46.

9. Wang, H., et al., Atlas of nonribosomal peptide and polyketide biosynthetic pathways reveals common occurrence of nonmodular enzymes. Proc Natl Acad Sci U S A, 2014. 111(25): p. 9259–64.

10. Schwarzer, D., R. Finking, and M.A. Marahiel, Nonribosomal peptides: from genes to products. Nat Prod Rep, 2003. 20(3): p. 275–87.

11. Sussmuth, R.D. and A. Mainz, Nonribosomal Peptide Synthesis-Principles and Prospects. Angew Chem Int Ed Engl, 2017. 56(14): p. 3770–3821.

12. Felnagle, E.A., et al., Nonribosomal peptide synthetases involved in the production of medically relevant natural products. Mol Pharm, 2008. 5(2): p. 191–211.

13. Drake, E.J., et al., Structures of two distinct conformations of holo-non-ribosomal peptide synthetases. Nature, 2016. 529(7585): p. 235–8.

14. Tanovic, A., et al., Crystal structure of the termination module of a nonribosomal peptide synthetase. Science, 2008. 321(5889): p. 659–63.

15. Beld, J., et al., The phosphopantetheinyl transferases: catalysis of a post-translational modification crucial for life. Nat Prod Rep, 2014. 31(1): p. 61–108.

16. Ciezki, K., et al., ngrA-dependent natural products are required for interspecies competition and virulence in the insect pathogenic bacterium Xenorhabdus szentirmaii. Microbiology (Reading), 2019. 165(5): p. 538–553.

17. Bushley, K.E. and B.G. Turgeon, Phylogenomics reveals subfamilies of fungal nonribosomal peptide synthetases and their evolutionary relationships. BMC Evol Biol, 2010. 10: p. 26.

18. Singh, S., et al., Role of secondary metabolites in establishment of the mutualistic partnership between Xenorhabdus nematophila and the entomopathogenic nematode Steinernema carpocapsae. Appl Environ Microbiol, 2015. 81(2): p. 754–64.

19. Roumia, A.F., Hussein, W., EI-Sayed, G. M., Elmasry, A. M. A., Fahim, S. F., In-Silco and In-Vitro Characterization of a Symbiotic Association Bacteria Isolated from Entomopathogenic Nematodes and Producers for Biological Control Non-Ribosomal Peptides. Egyptian Academic Journal of Biological Sciences, 2022. 14(1): p. 15.

20. Pantel, L., et al., Odilorhabdins, Antibacterial Agents that Cause Miscoding by Binding at a New Ribosomal Site. Mol Cell, 2018. 70(1): p. 83–94 e7.

21. Tobias, N.J., et al., Natural product diversity associated with the nematode symbionts Photorhabdus and Xenorhabdus. Nat Microbiol, 2017. 2(12): p. 1676–1685.

22. Crawford, J.M., et al., NRPS substrate promiscuity diversifies the xenematides. Org Lett, 2011. 13(19): p. 5144–7.

23. Davies, J., Specialized microbial metabolites: functions and origins. J Antibiot (Tokyo), 2013. 66(7): p. 361–4.

24. Demain, A.L. and A. Fang, The natural functions of secondary metabolites. Adv Biochem Eng Biotechnol, 2000. 69: p. 1–39.

25. Hibbing, M.E., et al., Bacterial competition: surviving and thriving in the microbial jungle. Nat Rev Microbiol, 2010. 8(1): p. 15–25.

26. Wang, Y., et al., Phenazine-1-carboxylic acid promotes bacterial biofilm development via ferrous iron acquisition. J Bacteriol, 2011. 193(14): p. 3606–17.

27. Dietrich, L.E., et al., The phenazine pyocyanin is a terminal signalling factor in the quorum sensing network of Pseudomonas aeruginosa. Mol Microbiol, 2006. 61(5): p. 1308–21.

28. Jiang, Q., et al., Quorum Sensing: A Prospective Therapeutic Target for Bacterial Diseases. Biomed Res Int, 2019. 2019: p. 2015978.

29. Zhao, X., Z. Yu, and T. Ding, Quorum-Sensing Regulation of Antimicrobial Resistance in Bacteria. Microorganisms, 2020. 8(3).

30. Kaplan, H.B. and E.P. Greenberg, Diffusion of autoinducer is involved in regulation of the Vibrio fischeri luminescence system. J Bacteriol, 1985. 163(3): p. 1210–4.

31. Seed, P.C., L. Passador, and B.H. Iglewski, Activation of the Pseudomonas aeruginosa lasI gene by LasR and the Pseudomonas autoinducer PAI: an autoinduction regulatory hierarchy. J Bacteriol, 1995. 177(3): p. 654–9.

32. Dadgostar, P., Antimicrobial Resistance: Implications and Costs. Infect Drug Resist, 2019. 12: p. 3903–3910.

33. Liedhegner, E., et al., Similarities in Virulence and Extended Spectrum Beta-Lactamase Gene Profiles among Cefotaxime-Resistant Escherichia coli Wastewater and Clinical Isolates. Antibiotics (Basel), 2022. 11(2).

34. Chen, S., et al., fastp: an ultra-fast all-in-one FASTQ preprocessor. Bioinformatics, 2018. 34(17): p. i884–i890.

35. Dobin, A., et al., STAR: ultrafast universal RNA-seq aligner. Bioinformatics, 2013. 29(1): p. 15–21.

36. Love, M.I., W. Huber, and S. Anders, Moderated estimation of fold change and dispersion for RNA-seq data with DESeq2. Genome Biol, 2014. 15(12): p. 550.

37. Jumper, J., et al., Highly accurate protein structure prediction with AlphaFold. Nature, 2021. 596(7873): p. 583–589.

38. Reuter, K., et al., Crystal structure of the surfactin synthetase-activating enzyme sfp: a prototype of the 4’-phosphopantetheinyl transferase superfamily. EMBO J, 1999. 18(23): p. 6823–31.

39. Pearson, W.R., An introduction to sequence similarity (“homology”) searching. Curr Protoc Bioinformatics, 2013. Chapter 3: p. 3 1 1–3 1 8.

40. Blin, K., et al., antiSMASH 7.0: new and improved predictions for detection, regulation, chemical structures and visualisation. Nucleic Acids Res, 2023. 51(W1): p. W46–W50.

41. Perry, R.D. and J.D. Fetherston, Yersiniabactin iron uptake: mechanisms and role in Yersinia pestis pathogenesis. Microbes Infect, 2011. 13(10): p. 808–17.

42. Dubos, R.J., Studies on a Bactericidal Agent Extracted from a Soil Bacillus : I. Preparation of the Agent. Its Activity in Vitro. J Exp Med, 1939. 70(1): p. 1–10.

43. Fuchs, S.W., et al., Structure elucidation and biosynthesis of lysine-rich cyclic peptides in Xenorhabdus nematophila. Org Biomol Chem, 2011. 9(9): p. 3130–2.

44. Fuchs, S.W., et al., Fabclavines: bioactive peptide-polyketide-polyamino hybrids from Xenorhabdus. Chembiochem, 2014. 15(4): p. 512–6.

45. Heinrich, A.K., et al., LuxS-dependent AI-2 production is not involved in global regulation of natural product biosynthesis in Photorhabdus and Xenorhabdus. PeerJ, 2017. 5: p. e3471.

46. Donmez Ozkan, H., et al., Nematode-Associated Bacteria: Production of Antimicrobial Agent as a Presumptive Nominee for Curing Endodontic Infections Caused by Enterococcus faecalis. Front Microbiol, 2019. 10: p. 2672.

47. Wenski, S.L., et al., Fabclavine diversity in Xenorhabdus bacteria. Beilstein J Org Chem, 2020. 16: p. 956–965.

48. Touray, M., et al., Natural products from Xenorhabdus and Photorhabdus show promise as biolarvicides against Aedes albopictus. Pest Manag Sci, 2024. 80(9): p. 4231–4242.

49. Pfeifer, B.A., et al., Biosynthesis of complex polyketides in a metabolically engineered strain of E. coli. Science, 2001. 291(5509): p. 1790–2.

50. Case, R.J., M. Labbate, and S. Kjelleberg, AHL-driven quorum-sensing circuits: their frequency and function among the Proteobacteria. ISME J, 2008. 2(4): p. 345–9.

51. Pflanze, S., et al., Nonribosomal peptides protect Pseudomonas nunensis 4A2e from amoebal and nematodal predation. Chem Sci, 2023. 14(41): p. 11573–11581.

52. Sun, S.J., et al., Cyclic Dipeptides Mediating Quorum Sensing and Their Biological Effects in Hypsizygus Marmoreus. Biomolecules, 2020. 10(2).

53. Wang, L., et al., luxS-dependent gene regulation in Escherichia coli K-12 revealed by genomic expression profiling. J Bacteriol, 2005. 187(24): p. 8350–60.

54. Marques, J.C., et al., LsrF, a coenzyme A-dependent thiolase, catalyzes the terminal step in processing the quorum sensing signal autoinducer-2. Proc Natl Acad Sci U S A, 2014. 111(39): p. 14235–40.

55. Marques, J.C., et al., Processing the interspecies quorum-sensing signal autoinducer-2 (AI-2): characterization of phospho-(S)-4,5-dihydroxy-2,3-pentanedione isomerization by LsrG protein. J Biol Chem, 2011. 286(20): p. 18331–43.

56. Ha, J.H., et al., Evidence of link between quorum sensing and sugar metabolism in Escherichia coli revealed via cocrystal structures of LsrK and HPr. Sci Adv, 2018. 4(6): p. eaar7063.

57. Medarametla, P., et al., Structural Characterization of LsrK as a Quorum Sensing Target and a Comparison between X-ray and Homology Models. J Chem Inf Model, 2021. 61(3): p. 1346–1353.

58. Miller, S.T., et al., Salmonella typhimurium recognizes a chemically distinct form of the bacterial quorum-sensing signal AI-2. Mol Cell, 2004. 15(5): p. 677–87.

59. Rutherford, S.T. and B.L. Bassler, Bacterial quorum sensing: its role in virulence and possibilities for its control. Cold Spring Harb Perspect Med, 2012. 2(11).

60. Zhen, X., et al., Economic burden of antibiotic resistance in ESKAPE organisms: a systematic review. Antimicrob Resist Infect Control, 2019. 8: p. 137.

61. Nelson, R.E., et al., National Estimates of Healthcare Costs Associated With Multidrug-Resistant Bacterial Infections Among Hospitalized Patients in the United States. Clin Infect Dis, 2021. 72(Suppl 1): p. S17–S26.

