## Supplemental Figures for "Quorum Sensing Regulators and Non-ribosomal Peptide Synthetases Govern Antibacterial Secretions in *Xenorhabdus szentirmaii*"

**Supplemental Figure 1: Analysis of 17 operons encoding NRPS modules in *X szentirmaii***

**Supplemental Figure 2: *X. szentirmaii* produces more potent antimicrobials than *X. nematophila***

**Supplemental Figure 3: The *lsr* operon in *X. szn c*ontains only LsrF and LsrG genes**

**Supplemental Table 1: RNA seq analysis**

**Supplemental Table 2: Up-regulated genes**

**Supplemental Table 3: NRPS genes**

**Supplemental Figure 1: Analysis of 17 operons encoding NRPS modules in *X szentirmaii***

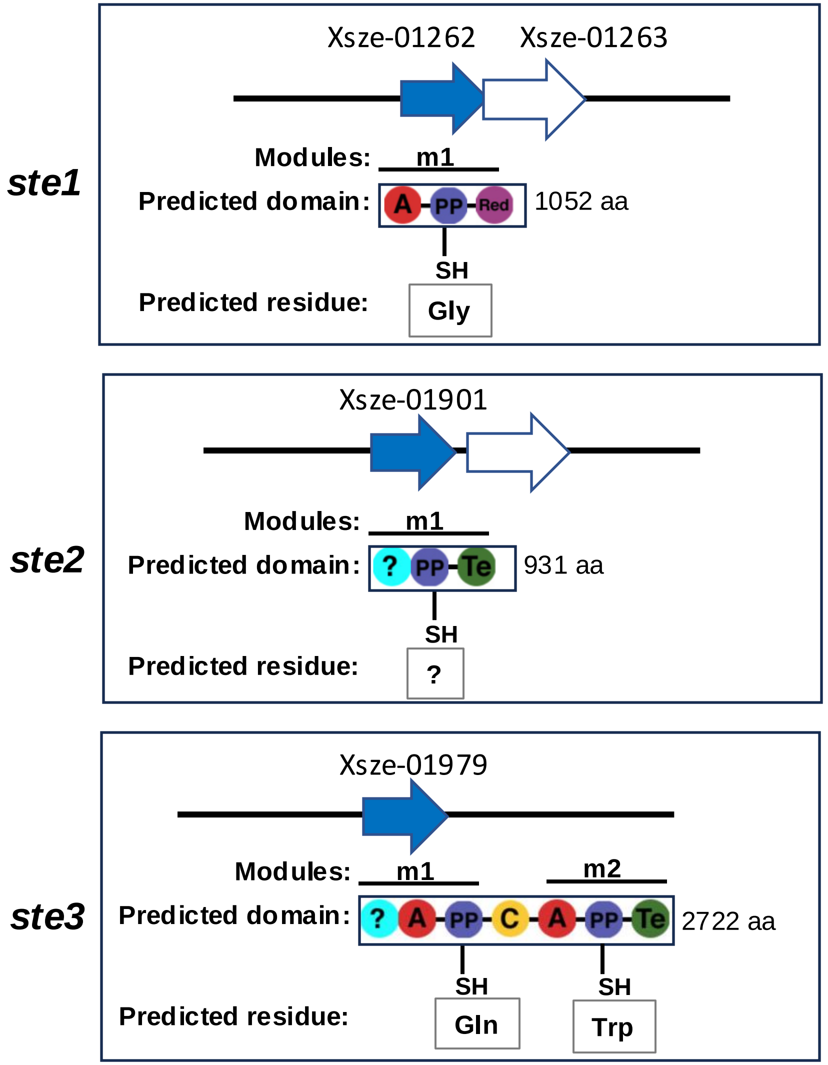


**Supplemental Figure 1:**

Upper panel: Schematic representation of operons ste1 to ste17, displaying the respective genes and its nomenclature (dark arrow) along with one or more than one adjacent gene (blank arrow).

Lower panel: Predicted NRPS modules from http://nrps.igs.umaryland.edu:

C = condensation domain,
A = adenylate domain,
PP =peptidyl carrier protein domain,
Te = Thioesterase domain,
? = unknow,
Cy = heterocyclization domain,
KS = ketosynthase domain,
KR = ketoreductase domain,
AT = acetyltransferase domain,
M = methyltransferase,
m = module,
aa = amino acids

**Supplemental Figure 1: Analysis of 17 operons encoding NRPS modules in *X szentirmaii***

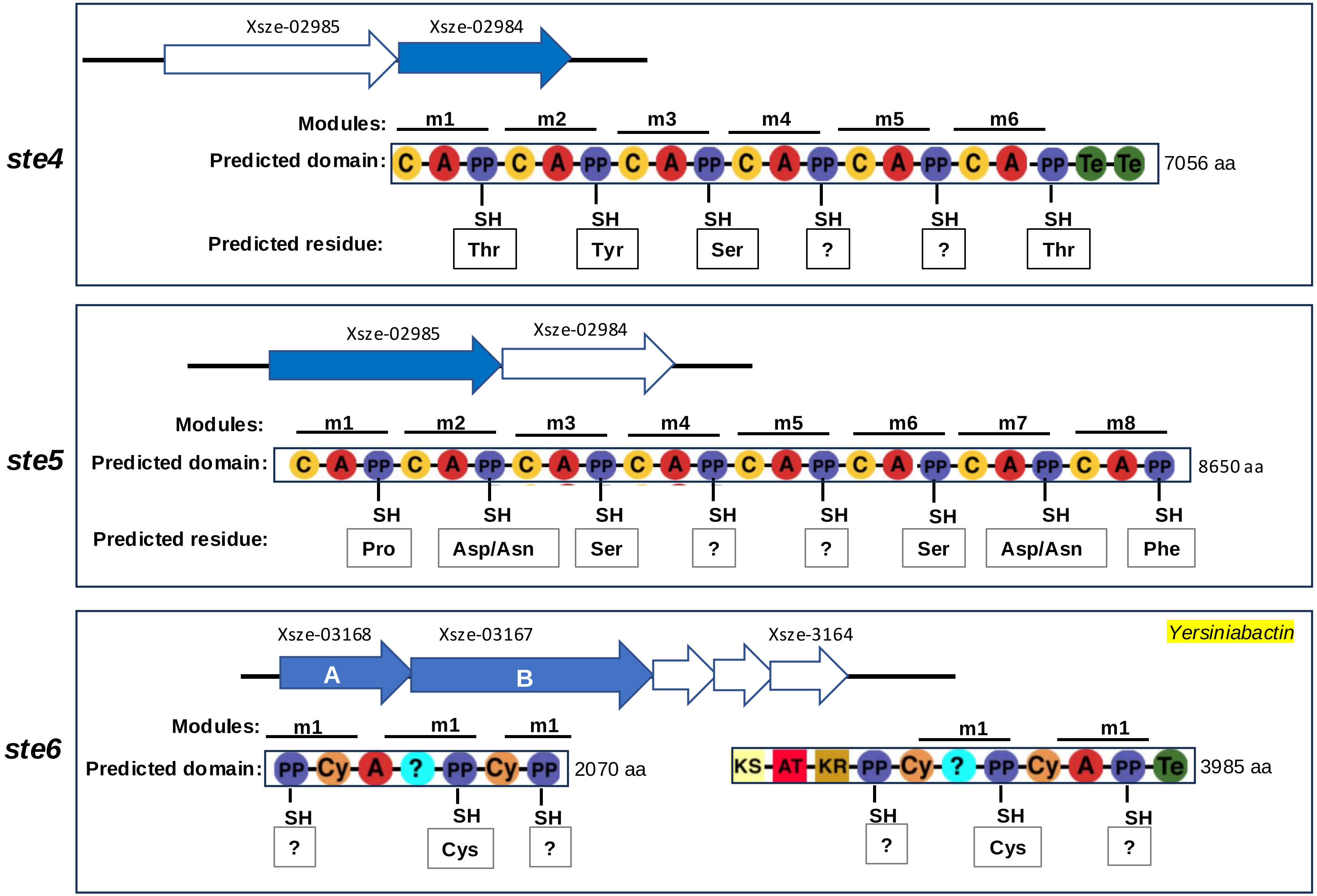

**Supplemental Figure 1: Analysis of 17 operons encoding NRPS modules in *X szentirmaii***

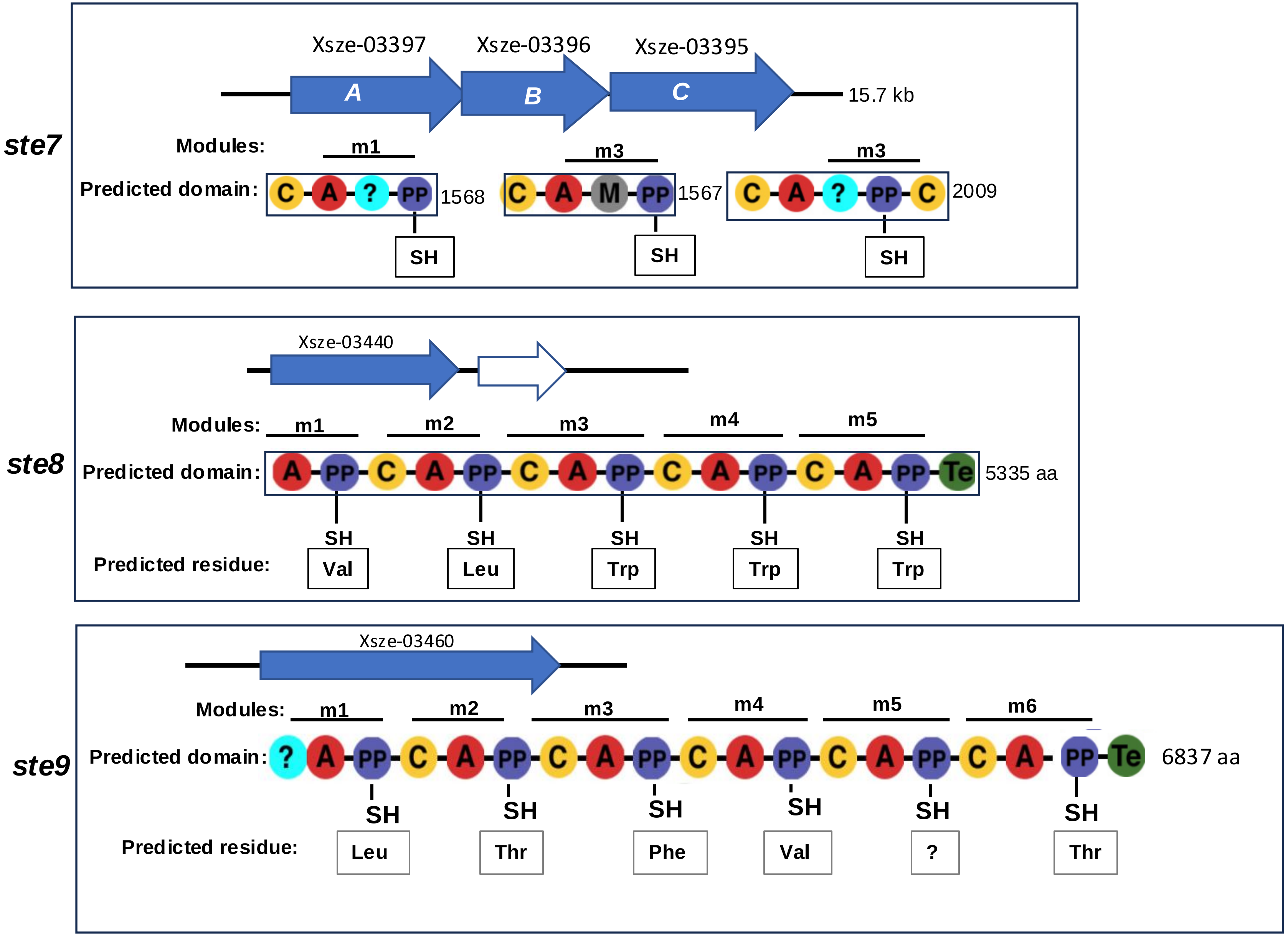

Supplemental Figure 1: Analysis of 17 operons encoding NRPS modules in *X szentirmaii*

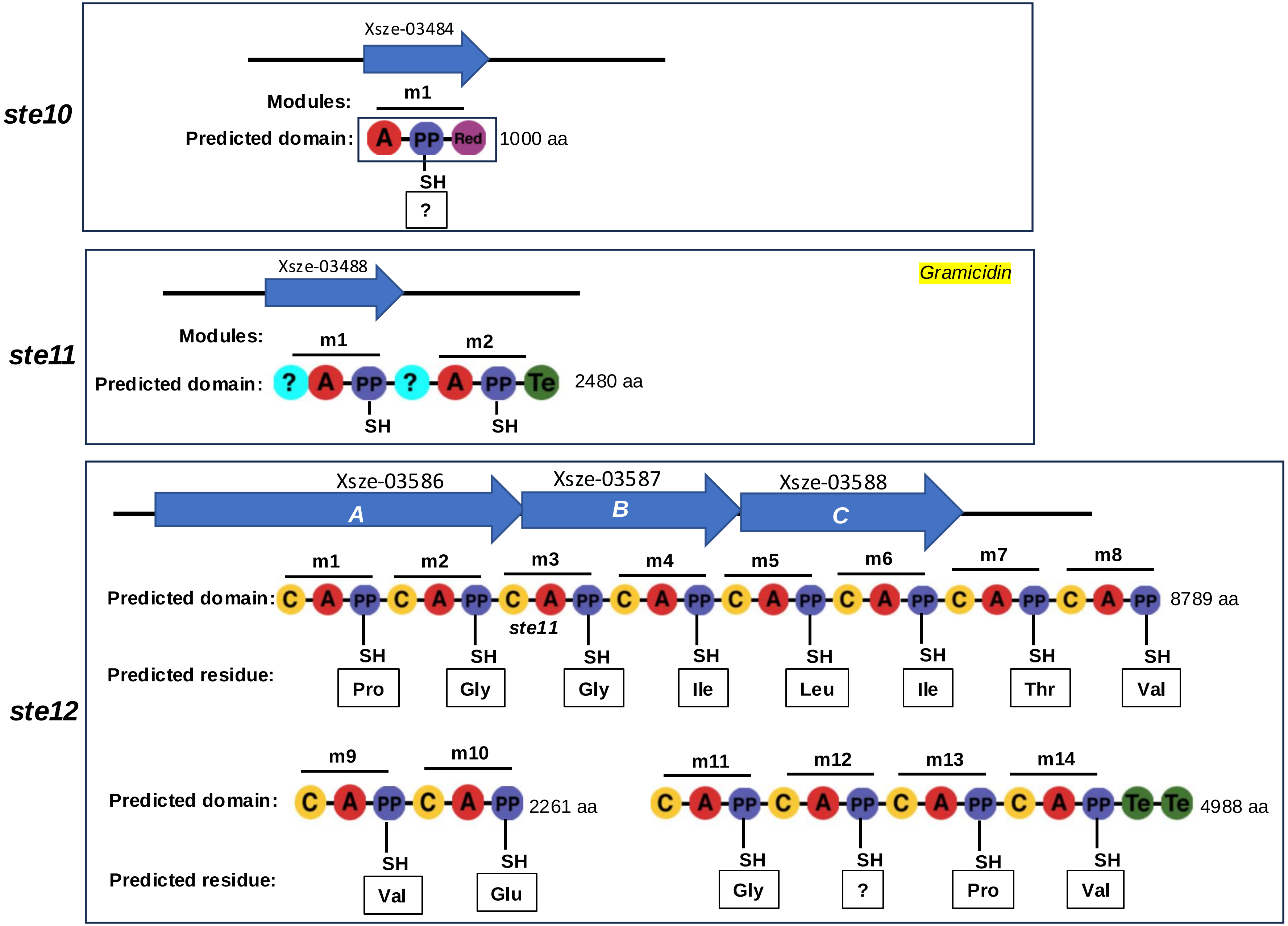

**Supplemental Figure 1: Analysis of 17 operons encoding NRPS modules in *X szentirmaii***

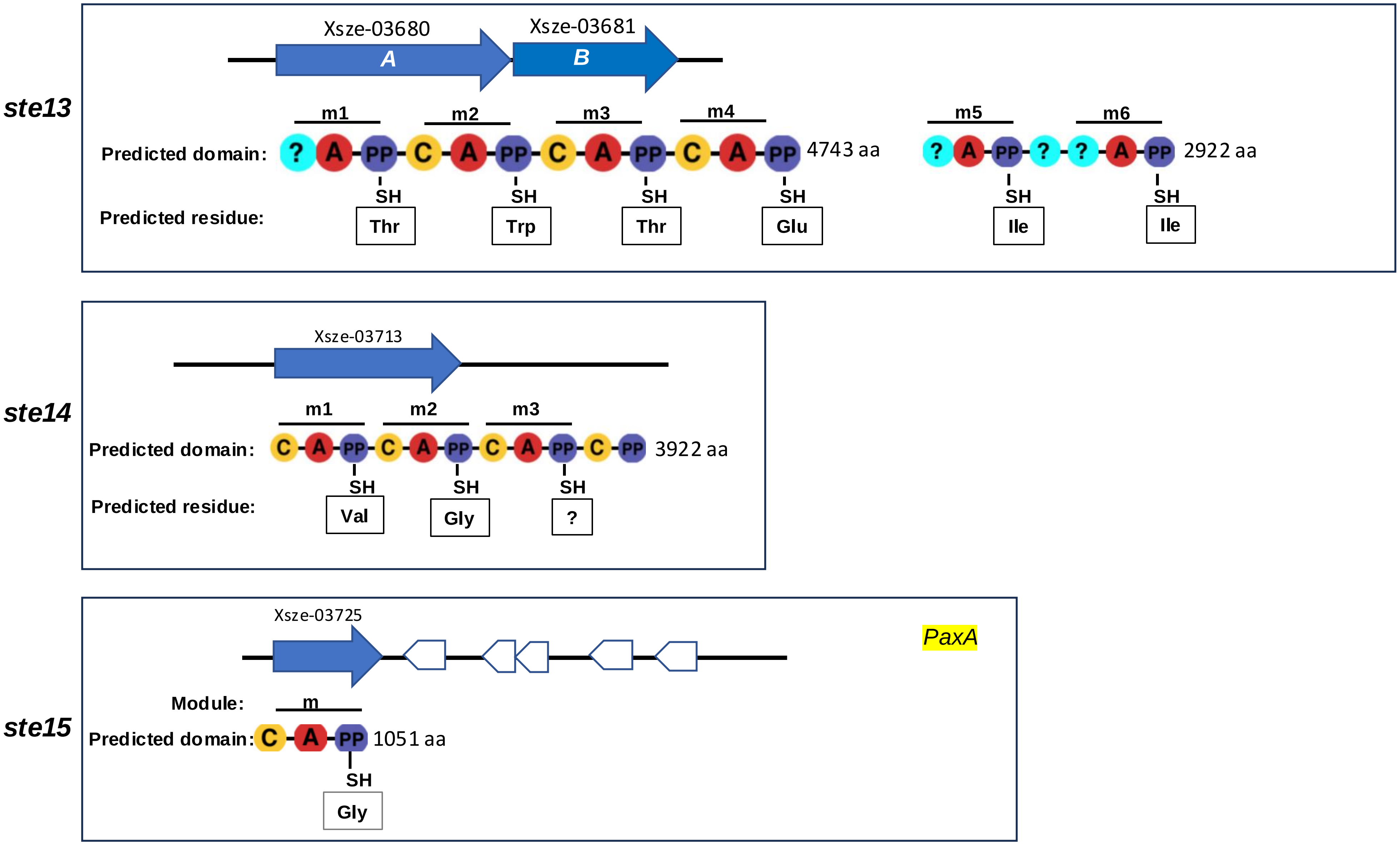

**Supplemental Figure 1: Analysis of 17 operons encoding NRPS modules in *X szentirmaii***

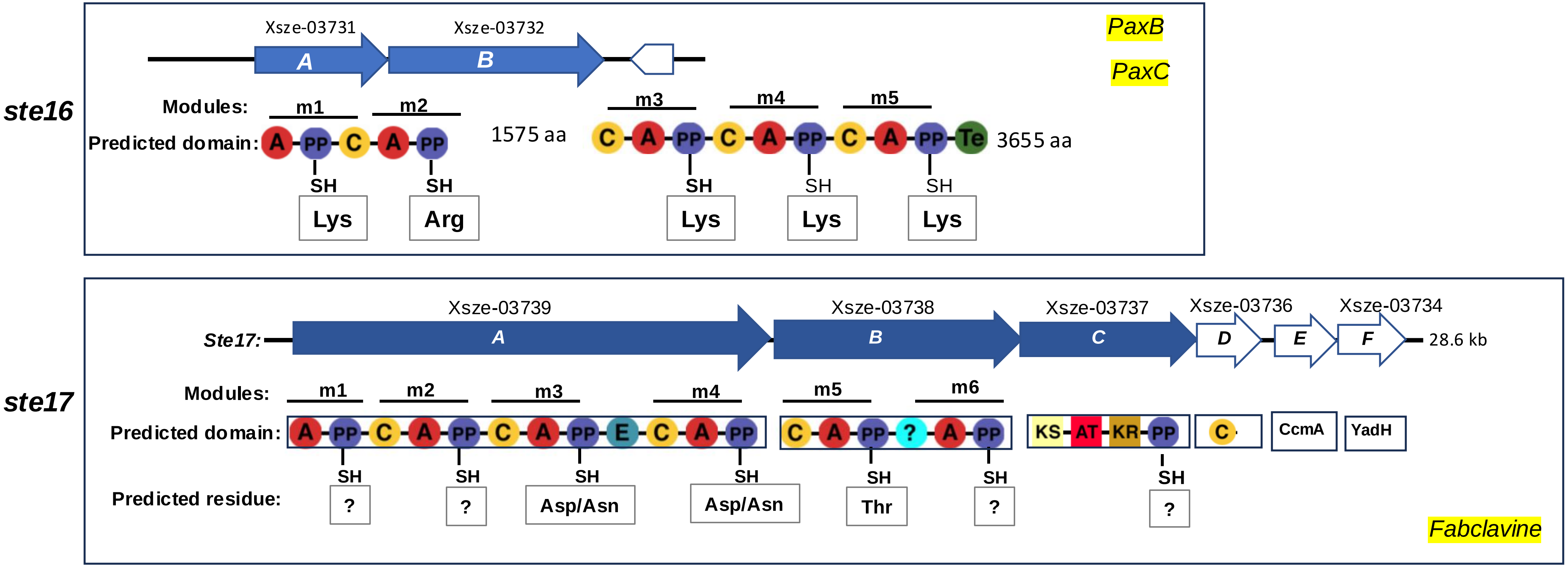

### Supplemental Figure 2: The *lsr* operon in *X szentirmaii* contains only LsrF and LsrG genes

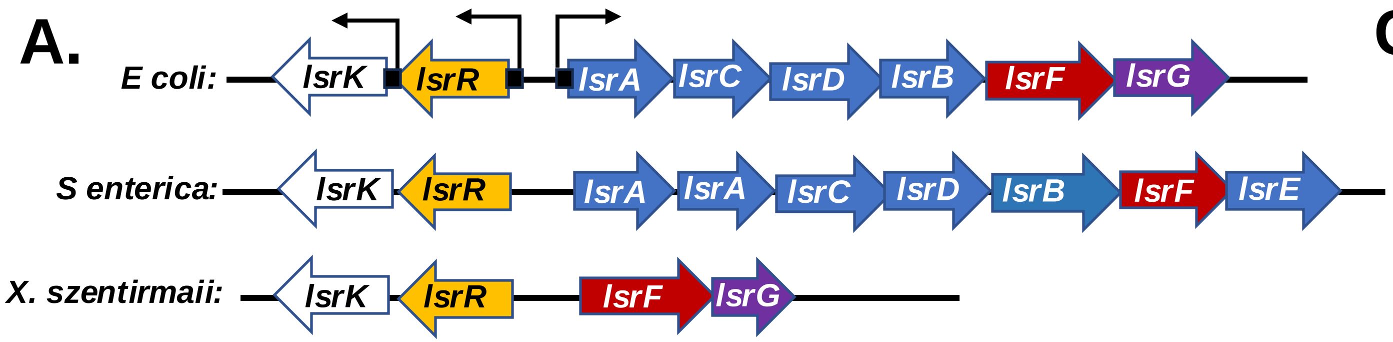


B.

```
E coli 1     MADLDDIKDGKDFRTDQPQQNIPFTLKGCGALDWGMQSRLSRIFNPKTGKTVMLAFDHGY 60
             MADLDDIKDGKDF  D PQ+N  F+LKGCGALDWGMQSRLSRIFNP T KTVMLAFDHGY
X szn  1     MADLDDIKDGKDFGIDTPQKNTLFSLKGCGALDWGMQSRLSRIFNPATNKTVMLAFDHGY 60

E coli 61    FQGPTTGLERIDINIAPLFEHADVLMCTRGILRSVVPPATNKPVVLRASGANSILAELSN 120
             FQGPTTGLERIDINIAPLFEH DVLMCTRGILRS VPPATNKPVVLRASGANSIL ELSN
X szn 61     FQGPTTGLERIDINIAPLFEHTDVLMCTRGILRSQVPPATNKPVVLRASGANSILTELSN 120

E coli 121   EAVALSMDDAVRLNSCAVAAQVYIGSEYEHQSIKNIIQLVDAGMKAGMPTMAVTGVGKDM 180
             EAVA++M+DA+RLN CAVAAQVYIGSE+EHQSIKNII+LVD GM+ GMPTMAVTGVGKDM
X szn  121   EAVAVAMEDALRLNVCAVAAQVYIGSEHEHQSIKNIIKLVDQGMRYGMPTMAVTGVGKDM 180

E coli 181   VRDQRYFSLATRIAAEMGAQIIKTYYVEKGFERIVAGCPVPIVIAGGKKLPEREALEMCW 240
              RDQRYFSLATRIAAEMGA IIKTYYV+ GFERI AGCPVPIVIAGGKKLPE EALEMC+
X szn  181   ARDQRYFSLATRIAAEMGANIIKTYYVDSGFERIAAGCPVPIVIAGGKKLPELEALEMCY 240

E coli 241   QAIDXGASGVDMGRNIFQSDHPVAMMKAVQAVVHHNETADRAYELYLSEK 290
             QAID GASGVDMGRNIFQS+ PVAM+KAVQAVVH NE   +AYEL+ +EK
X szn  241   QAIDQGASGVDMGRNIFQSESPVAMLKAVQAVVHQNEKPSQAYELFRTEK 290
```

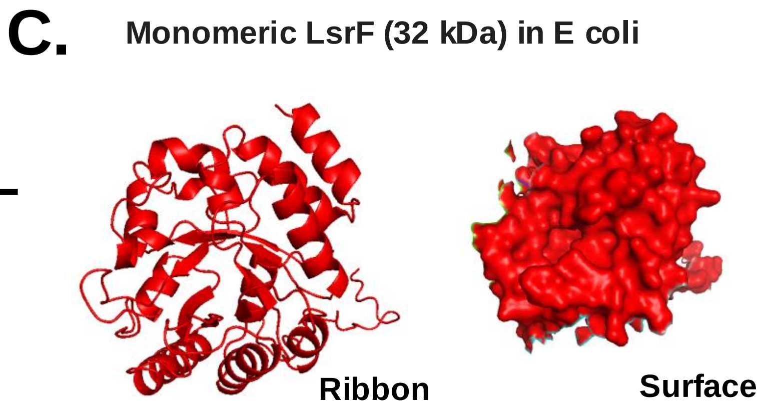


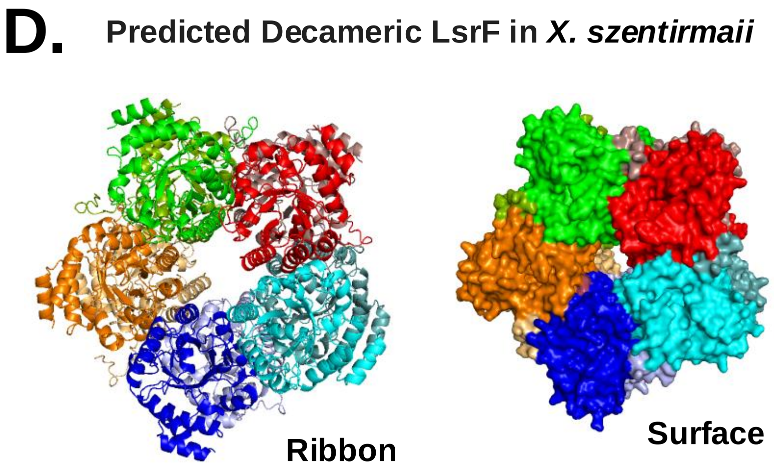


(A) Comparison of lsr operons among indicated bacterial species. Genes are indicated by arrows. (B) Comparative analysis of LsrF protein between E coli and *X szentirmaii* (X. szn). (C) Ribbon and surface diagram of the predicted decameric LsrF protein of *X szentirmaii* with each subunit represented in different color.
